# A global deep-sea small protein atlas reveals a reservoir of noncanonical antimicrobial peptides

**DOI:** 10.64898/2025.12.15.694180

**Authors:** Qiuyun Jiang, Yingchun Han, Chenyang Ye, Yiqian Duan, Fang Li, Zhuang Han, Célio Dias Santos-Júnior, Zhouqing Luo, Xiyang Dong

## Abstract

Small proteins encoded by small open reading frames (smORFs; ≤ 100 aa) represent a largely unexplored dimension of microbial diversity, especially in the deep sea. By analyzing 708 metagenomes from five major deep-sea biomes (hadal trenches, cold seeps, hydrothermal vents, abyssal plains, and seamounts), we constructed the Deep-Sea Small Protein Atlas, comprising 88.7 million smORFs with exceptional novelty and strong habitat specificity. Deep-learning predictions identified 5.47 million candidate antimicrobial peptides (c_AMPs), revealing a peptide space far larger and structurally distinct from known AMPs. Deep-sea c_AMPs are longer, enriched in nonpolar and acidic residues, and exhibit low charge and high intrinsic disorder, suggesting non-membranolytic modes of action. We synthesized 131 representative peptides, of which 87% were antimicrobial, with MICs as low as 1.25 μM, broad-spectrum antibacterial and even antifungal efficacy, and minimal mammalian cytotoxicity. Transcriptomics, TEM imaging, and peptide–protein modeling showed that representative peptides preserve membrane integrity while disrupting intracellular processes such as translation and metabolism, supporting intracellular, non-lytic mechanisms. This work uncovers a vast reservoir of previously unrecognized deep-sea small proteins and structurally unconventional AMPs, providing a foundational resource for discovering next-generation peptide therapeutics.

## Introduction

The deep sea, comprising cold seeps, hydrothermal vents, hadal trenches, abyssal plains, and seamounts, represents Earth’s largest and least explored biome, extending below 200 meters and encompassing nearly three-quarters of the ocean’s volume^1^. These environments are defined by extreme physicochemical conditions: perpetual darkness, low temperatures (0–4°C, except for hydrothermal systems that exceed 100°C), and immense hydrostatic pressure reaching up to 110 MPa in the deepest trenches. Such harsh conditions have driven the evolution of unique microbial communities with specialized physiological traits and metabolic architectures^1^. Deep-sea microorganisms are prolific producers of bioactive molecules that mediate competition and environmental adaptation, as exemplified by the discovery of unique biosynthetic gene clusters in cold-seep microbiome^2, 3^. Among these adaptations, small proteins encoded by small open reading frames (smORFs; ≤ 100 amino acids) have emerged as a particularly intriguing but underexplored molecular class^4^. Across diverse organisms, smORFs play critical roles in gene regulation^5^, stabilization of large protein complexes^6^, signal transduction^7^, protein transport^8^, stress responses^9^, and antimicrobial defense^10^. Despite this functional breadth, the diversity, ecological distribution, and biological roles of smORFs in deep-sea ecosystems remain largely unknown.

Antimicrobial peptides (AMPs) represent one biologically important subset of smORF-encoded proteins. These peptides exhibit activity against bacteria, fungi, and viruses^10, 11^, and are characterized by properties such as reduced propensity for resistance development^12, 13^, increased target specificity, and often lower cytotoxicity compared with conventional antibiotics. However, the systematic discovery of AMPs has been historically constrained by technical limitations, including biased biochemical enrichment, incomplete mass spectrometry reference databases^14^, and challenges in distinguishing genuine short coding sequences from spurious predictions^15^. Recent advances in ribosome profiling^16^, proteomics^17^, comparative genomics^18^, and artificial intelligence^19^ have transformed our ability to identify small proteins at scale. For example, the Global Microbial Small Protein Catalog (GMSC) contains nearly one billion non-redundant smORFs from 75 environments^4^, while the AMPSphere database reports 863,498 predicted AMPs, more than half of which show antimicrobial activity upon synthesis^20^. Similar large-scale efforts in the human gut^19^ and marine biofilms^21^ have further demonstrated the abundance and functional potential of environmental AMPs. Yet, despite representing Earth’s largest reservoir of microbial novelty^22, 23^, the deep sea remains conspicuously underrepresented in these global surveys. Most existing studies rely on cultivation-dependent approaches^24–28^, which capture only a small fraction of deep-sea microbial diversity, leaving the vast majority of deep-sea smORFs and AMP-like peptides unexplored.

AMPs operate through diverse mechanisms of action. Classical membrane-disruptive pathways involve electrostatic interactions between cationic peptides and anionic bacterial surface components such as lipopolysaccharides^29^, followed by membrane destabilization through barrel-stave, toroidal pore, or carpet-like mechanisms^30^. Increasing evidence, however, indicates that many AMPs act through non-membranolytic mechanisms^31^, including inhibition of cell-wall synthesis, replication, transcription, translation, or other essential intracellular processes^32–34^. Such mechanistic diversity expands the functional landscape of antimicrobial peptides and offers opportunities to develop next-generation therapeutics with improved specificity and reduced host toxicity^32^. Given the extreme physicochemical constraints of deep-sea environments, it remains unclear whether deep-sea-derived AMPs follow classical lytic paradigms or instead encode fundamentally distinct modes of action.

Here, we present the Deep-Sea Small Protein Atlas, a comprehensive resource that systematically maps smORFs and antimicrobial peptides across five major deep-sea biomes. By integrating large-scale metagenomic data with advanced computational annotation and deep-learning-based peptide prediction, we construct non-redundant, habitat-resolved smORF catalogs that uncover a vast and previously inaccessible diversity of microbial small proteins and previously undescribed AMP candidates. We further combine targeted peptide synthesis with antimicrobial assays, transcriptomics, structural modeling, and ultrastructural imaging to interrogate the biological activities and mechanistic potential of representative smORF-derived AMPs. Together, this atlas establishes a foundational framework for understanding small-protein-mediated adaptation in extreme environments and expands the structural and mechanistic repertoire available for next-generation peptide therapeutics.

## Results

### A vast and previously uncharted deep-sea small-protein landscape

We constructed a large-scale atlas of microbial smORFs from 708 metagenomic samples across five ecologically distinct deep-sea ecosystems **(Fig. 1a; Supplementary Table 1)**. Within each ecosystem, smORFs (≤ 100 aa) were predicted from assembled contigs using a unified workflow and subsequently dereplicated to generate five non-redundant habitat-specific catalogs (**Methods; Fig. 1b**). In total, we identified 88.7 million predicted smORFs, including 49.9 million from hadal trenches, 32.5 million from cold seeps, 3.8 million from hydrothermal vents, 0.8 million from abyssal plains, and 1.8 million from seamounts, representing the most comprehensive deep-sea small-protein resource available to date (**Supplementary Fig. 1; Supplementary Table 2**).

**Figure 1.**
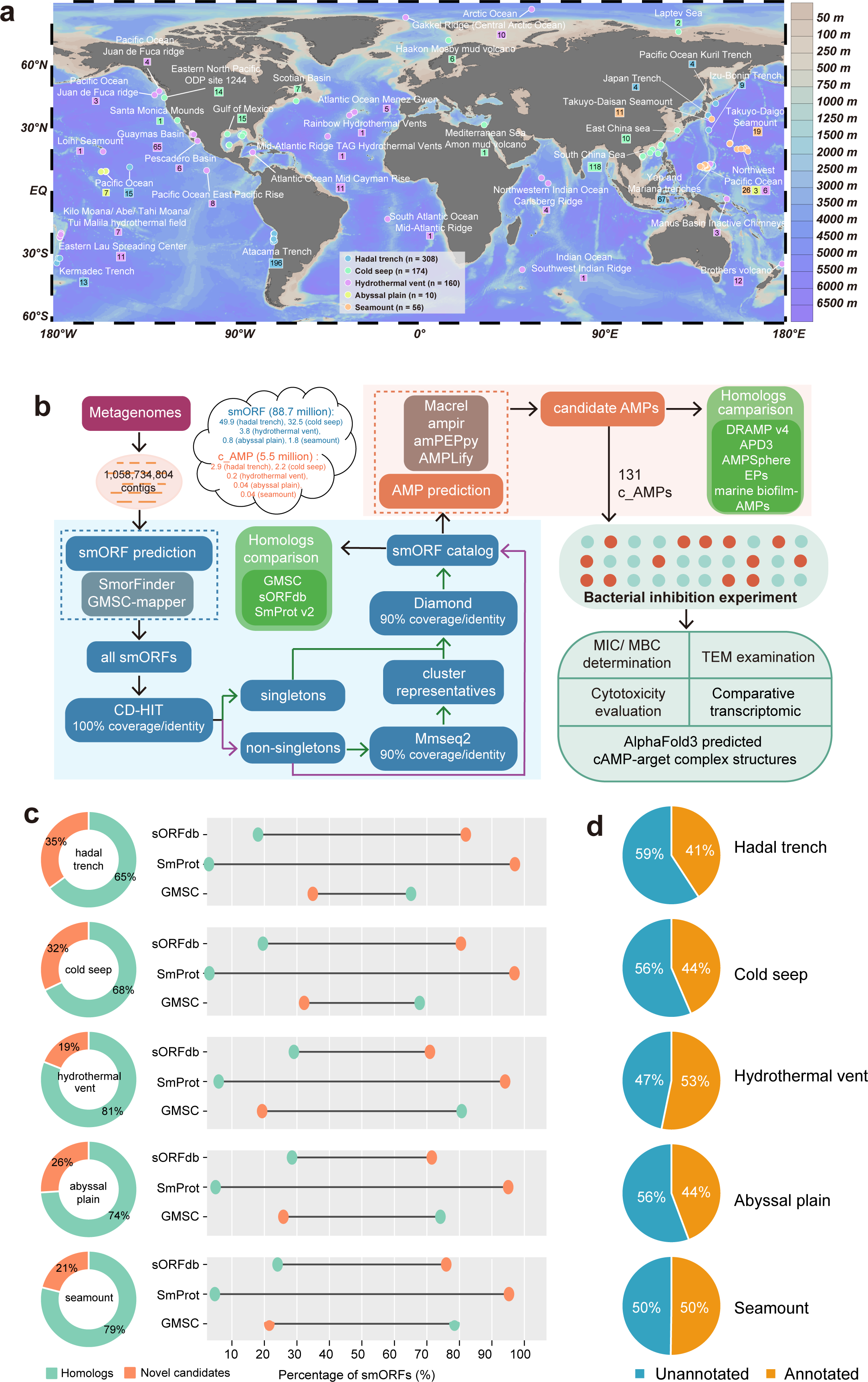
A global atlas of deep-sea small proteins with unprecedented diversity and novelty. **(a)** Geographic distribution of the deep-sea sampling sites included in this study. Circles denote metagenomic sampling locations, and squares indicate the number of samples collected at each site. Sampling and sequencing metadata for all datasets are provided in **Supplementary Table 1**. The world map was generated using Ocean Data View (ODV 5.1.7). **(b)** Workflow for predicting smORFs and candidate antimicrobial peptides (c_AMPs). Details are provided in methods. **(c)** Comparative homology landscape showing the novelty of deep-sea smORFs relative to existing small-protein databases. For each habitat, the proportions of smORFs with detectable homologs versus novel smORFs lacking database matches are shown. Some smORFs map to multiple databases; therefore, counts are not additive across databases. Detailed homology statistics are reported in **Supplementary Table 2**. **(d)** Functional annotation of smORF catalogs from the five deep-sea habitats, based on KEGG, eggNOG, CAZy, and Pfam databases. Summary statistics provided in **Supplementary Table 2**.

Deep-sea smORFs exhibited exceptionally high novelty relative to existing reference databases. On average, only 23.8% and 4.3% (range: 2.9–29.1%) of deep-sea smORFs displayed detectable homology to entries in sORFdb^35^ and SmProt^36^, respectively **(Fig. 1c)**. Even when compared with GMSC^4^, the largest microbial smORF dataset currently available, an average of 26.8% (range: 19.3–34.9%) of deep-sea smORFs remained unmatched. In total, 23.8 million smORFs showed no similarity to any tested database, greatly expanding the known sequence space of small proteins. Functional and taxonomic annotations further underscored this novelty. Approximately 53.6% (range: 47.8–59.2%) of smORFs lacked KEGG, eggNOG, CAZy, or Pfam annotations **(Fig. 1d; Supplementary Table 2)**, and nearly half (43.3%; range: 36.4–49.6%) could not be assigned to any taxonomic lineage across the five ecosystems (**Supplementary Fig. 2**). In line with estimates that ∼90% of deep-sea microbial taxa remain undescribed^22, 37^, the Deep-Sea Small Protein Atlas provides a foundational resource for uncovering previously unknown adaptive mechanisms, ecological functions, and bioactive molecules within Earth’s largest and least explored biosphere.

### Habitat specificity and distinct metabolic adaptations across deep-sea smORFs

Environmental heterogeneity across the five deep-sea biomes imposes strong selective pressures that shape the diversity and distribution of smORFs. Pairwise comparisons showed that smORFs are overwhelmingly habitat-specific: on average, 86.4% (62.9– 97.8%) of smORFs were restricted to a single biome, whereas only 4.4% (0.1–35.0%) were shared between any two habitats (**Fig. 2a; Supplementary Table 3**). Sharing was lowest between hydrothermal vents and abyssal plains (0.1%), reflecting the sharp ecological segregation associated with hydrothermal chemistry^38^. Consistently, within-habitat sharing was minimal, with only 2.5% of smORFs shared in hadal trench samples, 2.2% in cold seeps, 4.7% in hydrothermal vents, 37.1% in abyssal plains, and 21.2% in seamounts. The most physiochemically extreme habitats (hadal trenches, cold seeps, and hydrothermal vents) harbored the highest proportions of unique smORFs (> 95.3%), likely driven by steep redox gradients, high hydrostatic pressure, and strongly reducing conditions^38–41^. By contrast, the more stable, oxygenated abyssal plains and seamounts^42, 43^ exhibited comparatively higher levels of smORF sharing.

**Figure 2.**
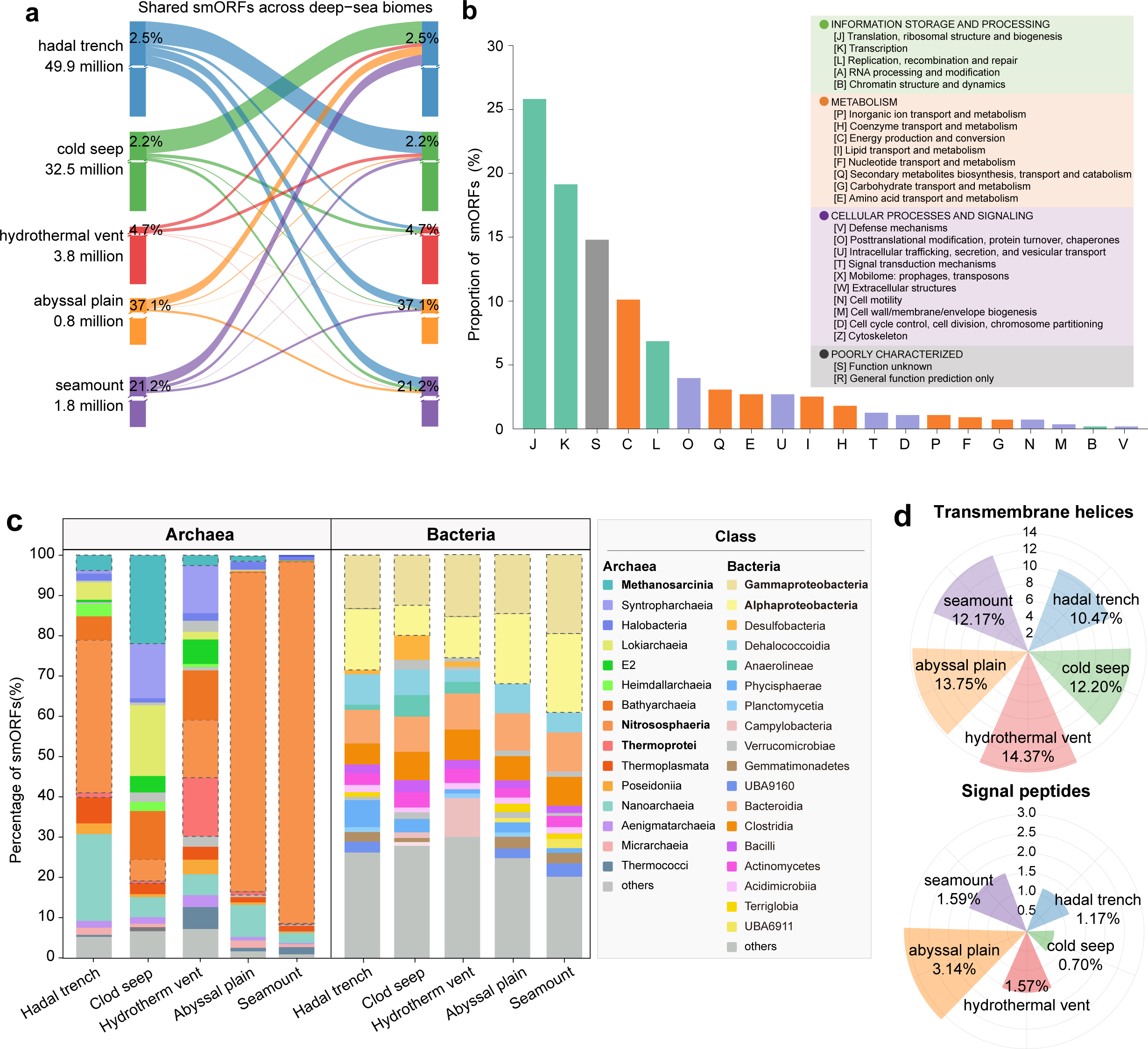
Habitat-specific patterns of deep-sea smORFs. **(a)** Shared smORFs across the five deep-sea habitats. Ribbon widths represent the proportion of smORFs shared between habitats, with numerical labels indicating the percentage of each habitat’s smORF catalog that is shared with others. Owing to the large disparities in smORF catalog sizes among habitats, truncated axes are used to improve visualization. Detailed pairwise comparisons are provided in **Supplementary Table 3**. **(b)** Functional annotation of shared small proteins based on the COG database. COG categories are grouped into four major functional classes. Colors correspond to individual COG categories within these groups. **(c)** Taxonomic annotation of smORFs across the five habitats. Stacked bar charts show the relative abundance of archaeal and bacterial taxa at class level. Taxonomic assignment was performed using GTDB release R220. Detailed habitat-specific taxonomic profiles are provided in **Supplementary Tables 6–7**. **(d)** Proportions of smORF-derived proteins predicted to contain transmembrane domains or signal peptides across habitats. Values shown in the radial plots indicate the percentage of small proteins carrying the corresponding structural feature in each habitat. Detailed information is provided in **Supplementary Table 9**.

Despite this pronounced habitat specialization, the atlas revealed a conserved core of 902 smORFs present across all biomes (**Supplementary Table 4**). Functional annotation showed enrichment in essential cellular processes, including translation, ribosomal structure and biogenesis (25.8%), transcription (19.1%), and energy production and conversion (10.1%) (**Fig. 2b; Supplementary Table 5**). These findings highlight a dual organizational structure within the deep-sea smORFome: a vast, highly specialized set of habitat-restricted smORFs shaped by localized selection pressures, and a small, deeply conserved core required for fundamental cellular processes across extreme physicochemical regimes.

Taxonomic profiling further underscored this pattern of habitat structuring while revealing remarkable phylogenetic breadth. Using MMseqs2-based classification^44^ with GTDB R220 ^45^, 46,303,778 smORFs were taxonomically assigned, spanning 18 archaeal and 169 bacterial phyla (**Supplementary Table 6**). Over half of all annotated smORFs were species-specific, with 81.8–89.7% of archaeal and 62.0–68.8% of bacterial assignments mapping to species-level lineages (**Supplementary Fig. 3**). Distinct habitat-level taxonomic signatures were evident: *Nitrososphaeria* dominated archaeal smORFs in hadal trenches (38.0%), abyssal plains (79.3%), and seamounts (89.9%), whereas *Methanosarcinia* (22.0%) and *Thermoprotei* (14.6%) predominated in cold seeps and hydrothermal vents, respectively (**Fig. 2c; Supplementary Table 7**). Among bacteria, *Alphaproteobacteria* were most abundant in hadal trenches (15.2%) and abyssal plains (17.4%), while *Gammaproteobacteria* predominated in cold seeps (12.5%), hydrothermal vents (15.5%), and seamounts (19.7%). These lineage-specific patterns are consistent with known biogeographic distributions^41, 43, 46^ and reflect microbial adaptation to highly contrasting redox, pressure, and energy landscapes. Collectively, the atlas demonstrates how local physicochemical regimes act as dominant selective forces shaping the evolution, diversification, and ecological partitioning of small proteins across the deep-sea biosphere^39, 47–49^.

Functional annotation further revealed metabolic specializations that distinguish deep-sea smORFs from global small-protein repertoires. On average, 46.4% of smORFs per habitat were assigned to COG categories^50^ (37.7 million total; **Supplementary Table 2**). Among these, ∼20.5% belonged to poorly characterized functions (S), highlighting the deep sea as a rich reservoir of uncharacterized protein functions. Annotated smORFs were enriched in categories C, J, K, E, and L, corresponding to energy metabolism, amino acid metabolism, and core informational processes (**Supplementary Figs. 4-5; Supplementary Table 8**). Consistent with GMSC database^4^, information-processing functions constituted a major fraction of annotated smORFs; however, the disproportionate representation of metabolic categories distinguishes the deep-sea smORFome from global trends^4^ and highlights the role of smORFs in mediating physiological adaptation to extreme environments^1, 3^.

Protein feature analysis showed that ∼12.6% of smORFs encode transmembrane helices and ∼1.6% contain signal peptides (**Fig. 2d; Supplementary Table 9**), suggesting that membrane-associated or secreted small proteins are widespread. These features were more common in bacterial than archaeal smORFs (**Supplementary Fig. 6**), likely reflecting both limited characterization of archaeal secretion systems and distinct ecological strategies. Approximately 1.1% of smORFs encoded Sec/Tat signal peptides and 0.5% encoded lipoprotein signal motifs **(Supplementary Fig. 7**), consistent with roles in cell–cell communication, nutrient acquisition, syntrophy, or environmental sensing^5, 51, 52^. Together, these findings reveal a deep-sea smORF repertoire enriched in metabolic and secreted functions that support microbial survival, interaction, and ecosystem functioning under extreme conditions.

### Unconventional, structurally flexible, and uncharacterized deep-sea c_AMPs

Using four independent deep-learning-based prediction tools, we identified 5,469,335 candidate antimicrobial peptides (c_AMPs) from the Deep-Sea Small Protein Atlas (**Supplementary Fig. 8**). This represents an expansion of ∼152-fold and ∼1,654-fold relative to the experimentally validated AMPs in DRAMP v4.0 ^53^ and APD v3 ^54^, respectively. The deep-sea repertoire also exceeds computationally predicted AMP collections by ∼6-fold (AMPSphere^20^), ∼127-fold (encrypted peptides (EPs) from the human proteome^55^), and ∼16,039-fold (marine biofilm-derived AMPs^21^). Sequence-level homology was minimal: only 0.03% and 0.21% of deep-sea c_AMPs showed similarity to DRAMP or APD entries, respectively (**Fig. 3a**). On average, 96.25% of c_AMPs were absent from all three major computational AMP resources (AMPSphere, EPs, and marine biofilms), and overall fewer than 14.32% had detectable homology to any known peptide. These results reveal a largely unexplored antimicrobial sequence space with substantial biotechnological potential.

**Figure 3.**
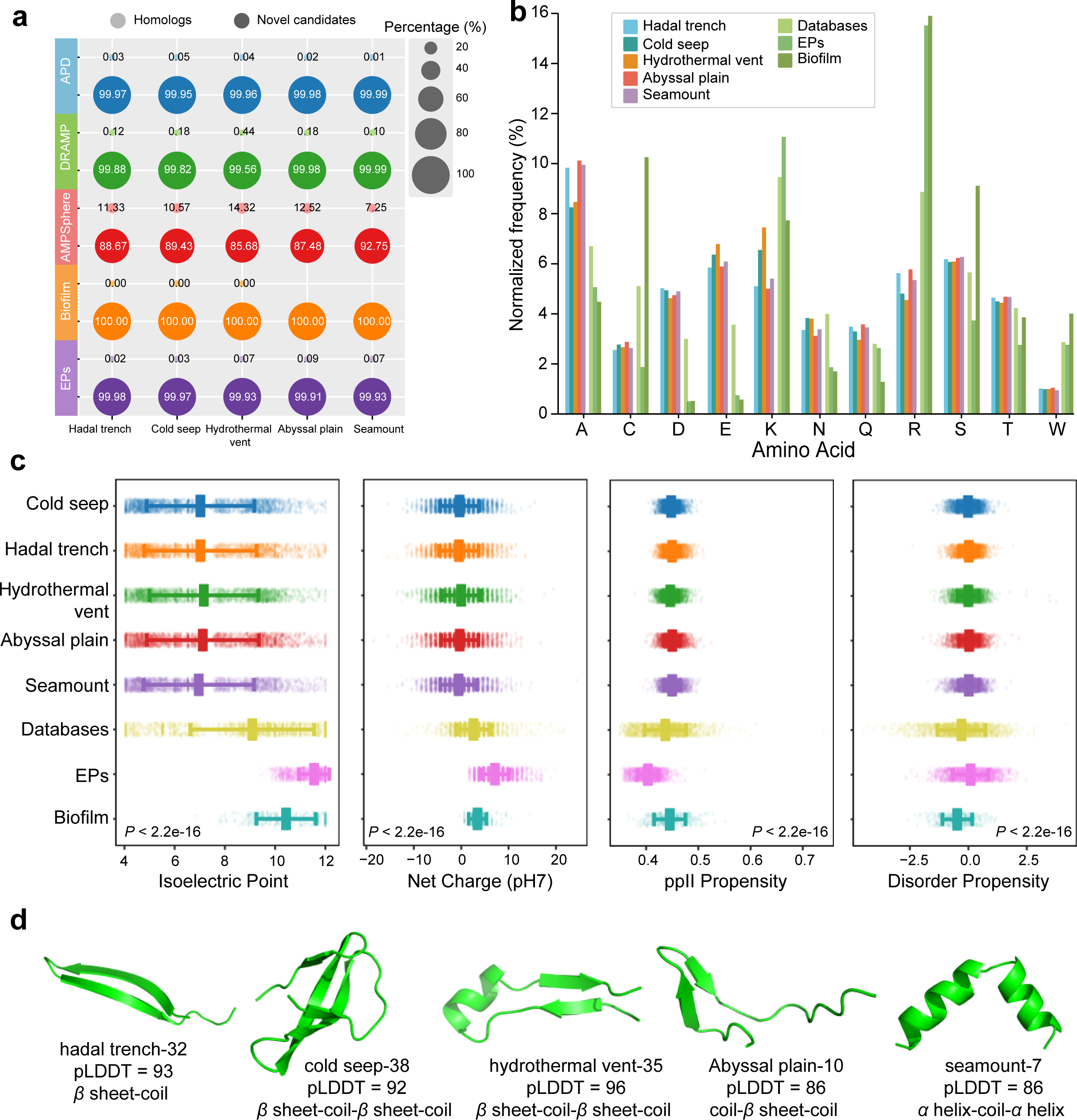
Defining a new structural and biophysical class of deep-sea antimicrobial peptides (c_AMPs). **(a)** Novelty assessment of c_AMPs identified across the five deep-sea habitats, benchmarked against experimentally validated AMP databases and large-scale predicted AMP resources. Bubble plots summarize the proportion of homologous (light color) versus novel sequences (dark color). **(b)** Normalized amino-acid composition profiles of deep-sea c_AMPs compared with reference AMP datasets. Detailed information is provided in **Supplementary Table 10**. **(c)** Multidimensional comparison of physicochemical descriptors across habitats and reference datasets. Detailed physicochemical properties and statistical data are provided in **Supplementary Tables 10-12. (d)** Representative AlphaFold2-predicted structures illustrating the conformational diversity of c_AMPs from different habitats. Model confidence is reported as per-residue pLDDT scores, highlighting ordered secondary-structure elements (*α*-helices, *β*-sheets) alongside flexible or disordered regions.

Amino acid composition profiling uncovered a distinct biochemical signature. Deep-sea c_AMPs were enriched in A (alanine), D (aspartic acid), E (glutamic acid), N (asparagine), Q (glutamine), S (serine), and T (threonine), but depleted in C (cysteine), K (lysine), R (arginine), and W (tryptophan) relative to DRAMP, APD, EPs and marine biofilm datasets (**Fig. 3b; Supplementary Table 10**). The enrichment of acidic residues (D, E) contrasts sharply with canonical membrane-disruptive AMPs, typically cationic and enriched in K and R that facilitate electrostatic interactions with negatively charged microbial membranes^20, 21, 55^, suggesting that these peptides likely employ fundamentally different interaction modes with microbial cells. Correspondingly, deep-sea c_AMPs exhibited near-neutral net charge (∼0 at pH 7) and substantially lower isoelectric points (pI ∼7) compared with reference AMPs (+2 to +7; pI 9–11.5, *P* < 0.0001; **Fig. 3c; Supplementary Tables 11–12**). Such electrostatic profiles imply a reduced dependence on charge-mediated membrane disruption, instead pointing toward alternative mechanisms, including metal ion coordination, protein–protein interaction, or intracellular pathway modulation^56–58^.

Multiple residue-level features further support a shift toward noncanonical structural paradigm. Elevated alanine content (**Fig. 3b**) suggests a bias toward flexible, low-complexity scaffolds^59^, whereas marked cysteine depletion reduces reliance on disulfide-stabilized *β*-sheet structures, favoring linear or intrinsically disordered conformations^60^. Predicted structural ensembles showed predominantly extended coils with short, transient *α*-helical segments (**Fig. 3d**), consistent with increased conformational plasticity. Such structural flexibility has been linked to non-membrane-disruptive antimicrobial modes, including selective intracellular targeting and specific molecular recognition^61, 62^. These observations align with our TEM imaging and transcriptomic analyses (see below), which showed antimicrobial activity without classical membrane permeabilization.

Physicochemical profiling reinforced the unique nature of deep-sea c_AMPs. They were longer (∼67 aa) and heavier (∼7 kDa) than reference AMPs (17–29 aa; 2–4 kDa), showed moderate hydrophobicity (GRAVY ≈ 0.01 vs. −0.30 to −0.47), and exhibited significantly reduced hydrophobic moments (∼0.38 vs. 0.72–0.77; *P* < 0.0001), indicative of weak *α*-helical amphipathicity (**Supplementary Fig. 9; Supplementary Tables 11–12**). Deep-sea peptides also displayed higher disorder propensity, enhanced polyproline II (PPII) tendency, and greater linear moment values (*P* < 0.05; **Fig. 3c; Supplementary Fig. 9**), features associated with increased structural flexibility, which may expand the repertoire of potential molecular targets. Reduced amphiphilicity and net charge could also decrease nonspecific cytotoxicity, a beneficial trait for therapeutic peptides. Together, these findings suggest that deep-sea c_AMPs represent a previously unrecognized class of antimicrobial peptides, characterized by low charge, intrinsic disorder, conformational adaptability, and diminished amphipathicity.

### Broad-spectrum antibacterial and antifungal activity with minimal cytotoxicity

To experimentally validate computational predictions in the Deep-Sea Small Protein Atlas, we synthesized 131 representative c_AMPs and evaluated their antimicrobial properties. Each peptide was screened at 125 μM, a benchmark concentration commonly used in AMP discovery studies^19^, against nine representative microorganisms, including Gram-positive bacteria (*Priestia filamentosa* and *Staphylococcus aureus* RN450), Gram-negative bacteria (*Escherichia coli* DH5α, *E. coli* BW25113, *Klebsiella pneumoniae* ATCC 13883, *Vibrio parahaemolyticus* VP.1997, *Halomonas bluephagenesis* TD01, and *Acinetobacter pittii*), and the yeast *Saccharomyces cerevisiae* BY4741. A total of 114 peptides (87.0%) inhibited the growth of at least one microorganism (**Supplementary Fig. 10; Supplementary Table 13**), demonstrating an exceptionally high validation rate compared with previous AMP discovery efforts.

Minimum inhibitory concentration (MIC) assays identified several highly potent peptides, including plain-4/sm-4, hydrothermal vent-19, and hydrothermal vent-35, with activities down to 1.25 μM, comparable to the positive controls nisin and melittin^63, 64^ (**Fig. 4a; Supplementary Tables 14–15**). Susceptibility varied across strains, with the highest numbers of active peptides observed for *P. filamentosa* (38 hits), *H. bluephagenesis* (31), *A. pittii* (16), and *S. aureus* (12). Several peptides displayed broad-spectrum properties, including 22/131 active against both Gram-positive and Gram-negative species and 7/131 active across multiple species within the same Gram group (**Supplementary Table 15**). Representative c_AMPs achieved complete bactericidal activity (≥ 99.9% killing) at 1 × – 4 × MIC against *P. filamentosa*, *H. bluephagenesis*, *V. parahaemolyticus*, *A. pittii*, and *E. coli* DH5*α* **(Fig. 4b; Supplementary Figs. 11– 13; Supplementary Table 16**), confirming strong and reproducible antimicrobial efficacy.

**Figure 4.**
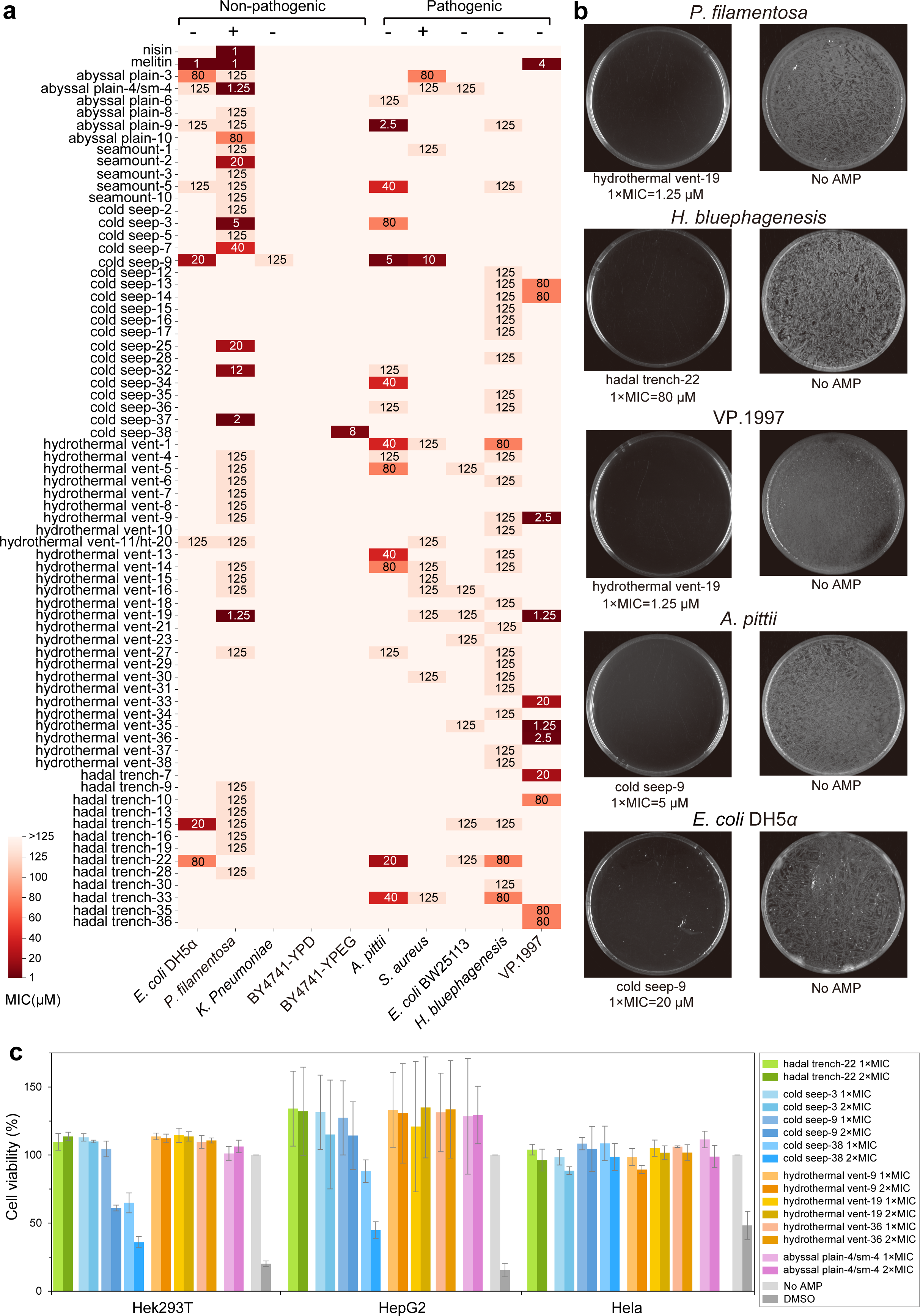
Functional validation of representative deep-sea c_AMPs. **(a)** Broad-spectrum inhibitory activity of synthetic c_AMPs against four non-pathogenic and five pathogenic microorganisms. MICs were determined using two-fold serial dilutions (2.5–125 μM) in 96-well plates containing ∼1 × 10^5^ CFU/mL. MIC values represent the mean concentration across three independent dilution series. Gram-positive (+) and Gram-negative (−) species are indicated. Detail information on MICs is provided in **Supplementary Tables 14-15**. **(b)** Bactericidal activity of representative c_AMPs at 1× MIC against *P. filamentosa*, *H. bluephagenesis*, *V. parahaemolyticus*, *A. pittii*, and *E. coli* DH5*α*. The minimum bactericidal concentration (MBC) was defined as the lowest concentration yielding ≥ 99.9% killing (≤ 0.1% survival). Representative plate images are shown. All assays were performed in triplicate. Detail information on MBCs is provided in **Supplementary Table 16**. **(c)** Cytotoxicity of synthetic c_AMPs toward mammalian cells (HEK293T, HepG2, HeLa) at 1–2 × MIC, assessed using standard metabolic-activity assays. Data represent mean ± SD from independent experiments. Detail data are provided in **Supplementary Table 17**.

Consistent with ecological expectations, no c_AMPs inhibited *S. cerevisiae* under fermentative conditions (YPD), potentially reflecting weaker evolutionary pressure to target fungi, an infrequent component of deep-sea communities (< 1–4%)^65, 66^. However, subtle growth defects emerged during the ethanol-dependent secondary growth phase (**Supplementary Fig. 14**). Under respiratory conditions (YPEG; ethanol + glycerol), cold seep-38 displayed strong antifungal activity (MIC = 8 μM) (**Fig. 4a**). Because ethanol metabolism requires functional mitochondria, these observations suggest that cold seep-38 may interfere with mitochondria-associated pathways, consistent with downstream transcriptomic responses described below.

To evaluate biosafety, we assessed eight deep-sea c_AMPs (hadal trench-22, cold seep-3, cold seep-9, cold seep-38, hydrothermal vent-9, hydrothermal vent-19, hydrothermal vent-36, and plain-4/sm-4) in mammalian cell lines, including a non-tumorigenic line (HEK293T) and two cancer lines (HepG2, HeLa). All peptides exhibited low cytotoxicity, with no significant dose-dependent effects observed in HeLa cells across the tested concentration range (**Fig. 4c; Supplementary Table 17**). The cold seep-9 showed mild dose-dependent toxicity toward HEK293T cells, whereas cold seep-38 displayed stronger, cell line–specific effects in HEK293T and HepG2 but minimal impact on HeLa cells. Given the differing energy demands and mitochondrial abundance across these cell types^67^, this cell line–specific response suggests that its activity may be linked to cellular energy metabolism and particularly mitochondria related function, consistent with the antifungal assays^68^.

### Diverse but predominantly non-lytic, intracellular antimicrobial strategies

Mechanistic profiling of representative deep-sea–derived c_AMPs revealed a coherent signature across peptides and taxa, most notably cold seep-9, hydrothermal vent-19, and cold seep-38, which showed the strongest *in vitro* activity against *A. pittii*, *P. filamentosa*, *V. parahaemolyticus*, and *S. cerevisiae*. Despite their structural diversity and the phylogenetic differences of the target organisms, these peptides consistently induced non-lytic antimicrobial responses characterized by modest perturbation of membrane permeability and pronounced intracellular effects on translation, proteostasis, and metabolic homeostasis, rather than catastrophic membrane lysis. To dissect these mechanisms, we performed transcriptomic profiling following AMP exposure, a widely used strategy for resolving intracellular targets and stress responses^31, 69^. Based on MIC results (**Fig. 4**), we selected four peptide–microbe combinations: *A. pittii* treated with cold seep-9 (MIC = 5 μM), *P. filamentosa* treated with hydrothermal vent-19 (MIC = 1.25 μM), *V. parahaemolyticus* treated with hydrothermal vent-19 (MIC = 1.25 μM), and *S. cerevisiae* treated with cold seep-38 (MIC = 8 μM). Across all organisms, only a restricted number of genes were significantly altered (|log2FC| ≥ 1 for bacteria, ≥ 0.5 for yeast; FDR < 0.05; **Fig. 5a**), consistent with targeted rather than global stress responses. DEG counts ranged from 127 in *A. pittii* to only 13 in *S. cerevisiae* (**Supplementary Tables 18-21**), reflecting organism- and pathway-specific peptide sensitivity.

**Figure 5.**
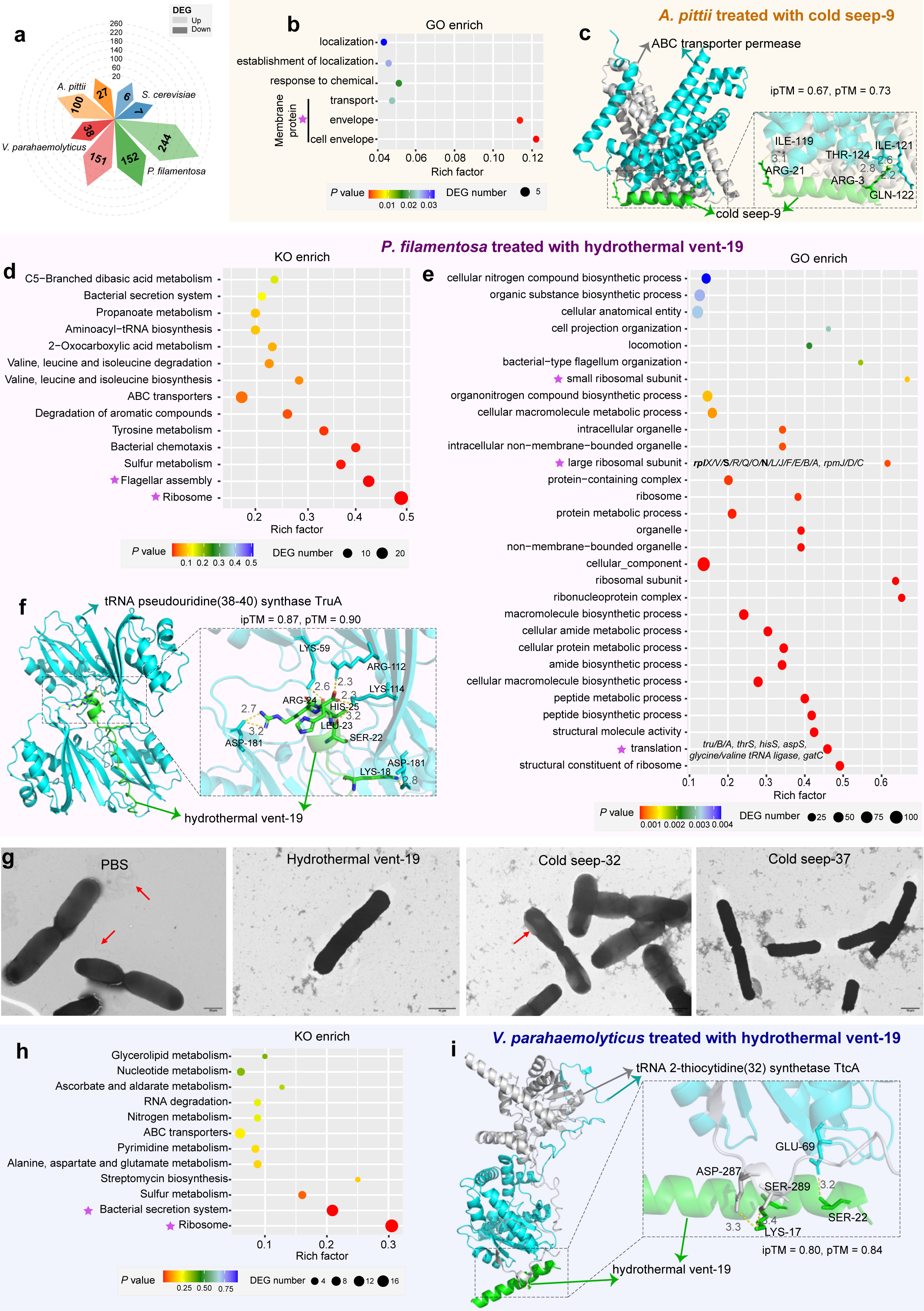
Mechanistic insights into deep-sea c_AMP activity across multiple microorganisms. **(a)** Diamond plots showing differentially expressed genes (DEGs) in four microorganisms treated with representative c_AMPs. Numbers within each sector indicate total DEGs, with light and dark colors marking significantly up- and downregulated genes, respectively. Detailed lists of up- and down-regulated genes for each treatment group are presented in **Supplementary Tables 18–21**. **(b–c)** Proposed mechanism of action of cold seep-9 in *A. pittii*, including GO enrichment analysis of DEGs (*P* < 0.05) (b) and a structural model of an ABC transporter permease subunit (light blue/gray) bound to cold seep-9 (green) (c). Protein–peptide complexes were generated using AlphaFold3, and model quality was assessed using ipTM and pTM scores. Insets highlight predicted interaction interfaces between the peptide and the permease. **(d–f)** Proposed mechanism of action of hydrothermal vent-19 in *P. filamentosa*, including KEGG KO enrichment analysis of DEGs (d), GO enrichment analysis (*P* < 0.005) (e), and an AlphaFold-based structural model of tRNA pseudouridine synthase TruA (light blue) bound to hydrothermal vent-19 (green) (f), with predicted peptide–protein interaction residues indicated. **(g)** Transmission electron microscopy (TEM) images of *P. filamentosa* treated with hydrothermal vent-19, cold seep-32, and cold seep-37 at 1× MIC. All TEM experiments were performed in triplicate. Scale bars: 10 μm. **(h–i)** Proposed mechanism of action of hydrothermal vent-19 in *V. parahaemolyticus*, including KEGG KO enrichment analysis showing significantly enriched pathways (h) and a structural model of tRNA 2-thiocytidine (32) synthetase (TtcA; light blue) bound to hydrothermal vent-19 (green), highlighting predicted interaction residues (i). All structural models were visualized using PyMOL.

In *A. pittii*, 120 of 127 DEGs were associated with metabolic pathways (**Supplementary Fig. 15a; Supplementary Table 18**). Carbon metabolism (21 DEGs) and amino acid metabolism (19 DEGs) were prominently affected. Two pathways showed the strongest perturbation: benzoate degradation (13/35 genes; **Supplementary Fig. 15a, 16b**) and branched-chain amino acid degradation (9/28 genes; **Supplementary Fig. 15a, 16c**). GO enrichment **(Fig. 5b)** highlighted a strong overrepresentation of membrane-localized proteins, particularly those associated with the cell envelope and transport processes. The transporters comprised systems mediating the uptake or export of carbon sources (e.g., *quiX*), amino acids (including multiple MFS and DMT family transporters), and ions (e.g., members of the RcnB family (**Supplementary Table 18**). Although cold seep-9 lacks predicted transmembrane helices (**Supplementary Fig. 16d**), AlphaFold3 modeling suggested a physically plausible interaction with an ABC transporter permease (ipTM = 0.67, pTM = 0.73; **Fig. 5c**), pointing to a mechanism involving modulation of membrane-associated transport rather than membrane insertion. Collectively, these findings suggest that cold seep-9 modestly perturbs membrane permeability, leading to disruption of carbon, nitrogen, and ion homeostasis and broadly impairing metabolic balance.

In *P. filamentosa*, translation-related genes were strongly downregulated (28/57 ribosomal genes), alongside genes involved in flagellar assembly (15/35) (**Fig. 5d; Supplementary Fig. 15b; Supplementary Table 19**). Both large and small ribosomal subunits exhibited extensive downregulation (**Fig. 5e**). Aminoacyl-tRNA ligases (*thrS* etc), tRNA-modifying enzymes (including *truA* and *truB*), and aminotransferases (*gatC*) were also repressed (**Supplementary Table 19**), indicating substantial impairment of translational capacity. AlphaFold3 modeling predicted a high-confidence interaction between hydrothermal vent-19 and TruA (ipTM = 0.87, pTM = 0.90), a key tRNA pseudouridine synthase (**Fig. 5f**), providing a mechanistic basis for translation inhibition. Altogether, these results suggest that hydrothermal vent-19 likely exerts its antibacterial activity primarily by disrupting protein translation, thereby impairing multiple cellular processes including flagellum biogenesis, ultimately compromising bacterial viability. Consistent with the well-documented flagellar shedding phenotype observed following AMP treatment^70^, TEM imaging revealed flagellar loss in hydrothermal vent-19–treated cells (**Fig. 5g**). Additional AMPs exhibited distinct phenotypes: cold seep-32 caused cell enlargement with pale cytoplasmic density, whereas cold seep-37 induced elongation and cytoplasmic leakage, highlighting mechanistic diversity among deep-sea peptides.

In *V. parahaemolyticus*, hydrothermal vent-19 triggered upregulation of ribosomal genes (17/56) and components of the bacterial secretion system (12/58) (**Fig. 5h; Supplementary Fig. 14c; Supplementary Table 20**), critical for virulence^71^. This response contrasts with the downregulation observed in Gram-positive *P. filamentosa*, suggesting that organism-specific regulatory architectures shape transcriptional outputs despite potentially shared molecular targets. Supporting this, hydrothermal vent-19 also displayed a predicted interaction with TtcA (ipTM = 0.8, pTM = 0.84; **Fig. 5i**), a tRNA thiolation enzyme involved in tRNA modification that contributes to efficient translation, particularly under stress conditions. Together, these data indicate that hydrothermal vent-19 primarily targets translation machinery, with downstream consequences that differ between Gram-positive and Gram-negative bacteria.

The limited DEGs in *S. cerevisiae* (n = 13; |log2FC| ≥ 0.5, FDR < 0.05; **Supplementary Fig. 15d; Supplementary Table 21)** is consistent with the observation that cold seep-38 treatment resulted in only slight growth reduction during the secondary growth phase in YPD, when ethanol becomes the primary carbon source **(Supplementary Figs. 14, 15d)**. The complete growth inhibition under YPGE condition **(Fig. 4a)** prompted us to further investigate potential transcriptional changes, revealing that cold seep-38 induces mild rather than extensive transcriptional perturbation. KEGG enrichment revealed perturbation of carbon and energy metabolism, with *PCK1*, a key gluconeogenic and ethanol-utilization gene (**Supplementary Figs. 15d, 17**), significantly downregulated. Removal of the peptide fully restored respiratory growth (**Supplementary Fig. 18**), indicating a reversible inhibitory mechanism. The peptide’s cell-line–specific cytotoxicity (**Fig. 4c**) further supports a connection to mitochondrial or energy-linked processes, raising the possibility that cold seep-38 may function as a reversible modulator of central carbon metabolism with broader implications for immunometabolism and cancer biology (**Supplementary Fig. 19**).

## Discussion

This study provides three conceptual advances: (i) the first global atlas of deep-sea small proteins; (ii) the discovery of a structurally distinct class of noncanonical c_AMPs; and (iii) experimental evidence that representative peptides act predominantly through intracellular, non-lytic mechanisms. The Deep-Sea Small Protein Atlas uncovers an expansive and previously inaccessible repertoire of bioactive peptides, over 85% of which show no detectable similarity to existing AMP databases, greatly expanding the known sequence space of natural antimicrobials. Representative c_AMPs demonstrated broad antimicrobial efficacy across Gram-positive and Gram-negative bacteria and even a mitochondrial-dependent yeast strain, suggesting that their primary targets may lie beyond the cell membrane. Such breadth of activity is consistent with mechanisms that are not strictly dependent on membrane composition^32^. Indeed, transcriptomic analyses revealed perturbations in translation, proteostasis, and metabolic pathways rather than gene expression patterns characteristic of classical membrane lysis, indicating that intracellular pathways are likely the dominant targets of these peptides under deep-sea environmental conditions such as high hydrostatic pressure, extreme temperature, and nutrient limitation.

A distinctive feature of the deep-sea c_AMPs is their predicted intrinsic disorder. Unlike canonical AMPs that fold into amphipathic *α*-helices, these peptides appear structurally plastic, favoring extended or low-complexity conformations. Intrinsically disordered peptides can undergo environment-dependent structural transitions^72^ and may interact with membranes through transient, peripheral contacts rather than inserting into bilayers to form stable pores. The behavior of cold seep-9 exemplifies this: despite lacking predicted transmembrane helices, it alters membrane permeability sufficiently to disrupt carbon, nitrogen, and ion homeostasis while maintaining overall membrane integrity. Analogous to disordered C-terminal motifs in endocytic adaptors, which generate steric pressure to drive membrane curvature^73^, deep-sea peptides may exert functional effects without requiring stable secondary structure.

Functionally, representative c_AMPs inhibited diverse microorganisms, yet TEM imaging revealed no evidence of catastrophic membrane damage. Cells maintained intact envelopes but exhibited altered intracellular organization and transcriptional responses. These observations contrast sharply with membrane-disruptive AMPs, such as polymyxins and many amphipathic helical peptides^19, 20^, which kill by compromising membrane integrity. While membrane lysis is effective, it often correlates with cytotoxicity in mammalian cells^74^ due to nonspecific interactions with eukaryotic membranes. In contrast, several antimicrobial peptides can cross membranes without causing detectable damage and modulate intracellular processes^32^, an emerging but still poorly understood antimicrobial strategy. The deep-sea peptides characterized here appear to fall within this underexplored mechanistic class.

Together, these findings reveal a vast diversity of previously uncharacterized small proteins encoded by extreme marine ecosystems. The c_AMPs uncovered here possess a unique combination of low net charge, intrinsic disorder, structural plasticity, and intracellular, non-lytic mechanisms of action, collectively expanding the conceptual landscape of natural antimicrobial strategies. These results challenge the prevailing emphasis on membrane-disruptive paradigms and suggest that environmental extremes such as those in the deep sea may drive the evolution of fundamentally distinct antimicrobial modalities.

At the same time, our mechanistic analyses were restricted to a subset of short, synthesizable peptides, and it remains unknown whether longer or more complex c_AMPs share similar structural or mechanistic properties. Future work will require pinpointing precise intracellular targets, determining peptide–protein interaction structures at atomic resolution, exploring functional diversity across peptide families, and evaluating pharmacokinetics and therapeutic potential *in vivo*. By enabling systematic discovery and mechanistic exploration, the Deep-Sea Small Protein Atlas provides a foundational resource for identifying next-generation peptide antibiotics inspired by Earth’s most remote and evolutionarily unique environments.

## Materials and Methods

### Metagenomic dataset and processing

Metagenomic datasets included 308 samples (40 water samples and 268 sediment samples) from hadal trenches in the Pacific Ocean (Mariana, Kermadec, Atacama, Kuril, Izu-Bonin, and Japan Trenches), 174 cold seep sediment samples collected from 17 globally distributed seep sites, 160 hydrothermal vent samples (three hydrothermal chimney samples, seven rock samples, 28 microbial mat samples, 38 water samples, and 84 sediment samples) from the Pacific, Atlantic, Indian, and Arctic Oceans (Brothers Volcano, Guaymas Basin, Juan de Fuca Ridge, Gakkel Ridge, Manus Basin, the Southern Mariana Trough, and the Mid Cayman Rise), 56 seamount sediment samples from the Pacific Ocean (Northwest Pacific Ocean, Takuyo-Daigo Seamount, Takuyo-Daisan Seamount, and seamounts along the Yap and Mariana trenches), and 10 abyssal plain sediment samples from the Pacific Ocean **(Supplementary Fig. 1a and Supplementary Table 1)**. A total of 115 cold seep sediment metagenomic datasets were generated by our group and reported in our previous publications^23, 75–82^, whereas the remaining datasets were retrieved from the National Center for Biotechnology Information Sequence Read Archive (NCBI-SRA) and the European Bioinformatics Institute European Nucleotide Archive (EBI-ENA). Detailed information, including associated publications, sample metadata, sequencing platforms, and accession numbers, is provided in **Supplementary Table 1**. For each sample, metagenomic reads were quality-filtered using fastp^83^ (v0.23.2, default parameters), and clean reads were assembled using MEGAHIT^84^ (v1.2.9, default parameters). In total, 1,058,734,804 contigs were generated (> 200 bp), including 807,712,620 from hadal trenches, 59,011,641 from cold seeps, 125,126,723 from hydrothermal vents, 15,953,963 from abyssal plains, and 50,929,857 from seamounts (**Fig. 1b and Supplementary Table 2**). All assembled contigs were used for subsequent smORF prediction.

### Non-redundant smORF catalog construction and cross-habitat analyses

smORFs were predicted from assembled contigs using smorFinder^85^ (v1.0.0, default parameters) and GMSC-mapper^4^ (v0.0.1a0, parameters: --no-habitat --no-taxonomy). The predicted smORFs were first deduplicated at 100% amino acid identity and 100% coverage using CD-HIT^86^ (v4.8.1, parameters: -c 1 -aS 1 -g 1 -d 0). All non-singleton sequences were subsequently clustered at 90% amino acid identity and coverage with MMseqs2 (v13.45111)^87^ using the easy-linclust module (parameters: -c 0.9 -min-seq-id 0.9). Singletons removed during the 100% identity deduplication step were further evaluated to retain potentially meaningful homologs. These sequences were aligned against MMseqs2 cluster representatives using DIAMOND^88^ (v2.1.8, parameters: --id 90 --query-cover 90 --subject-cover 90 -e 1e-5). Homologous singletons were merged with the non-singleton sequences identified earlier to generate the final non-redundant smORF catalogs for each deep-sea habitat. The overall workflow is summarized in **Fig. 1b**. To assess smORF sharing across habitats, pairwise comparisons between smORF catalogs were performed using DIAMOND (v2.1.8, parameters: --id 100 --query-cover 100 --max-target-seqs 1). smORFs with hits meeting this criterion were considered shared.

### Taxonomic and functional annotation of smORFs

Taxonomic assignment of smORFs was performed using MMseqs2 ^44^ (v13.45111, taxonomy module; parameter: --tax-lineage 1) against the GTDB reference database^45^ (R220). Functional annotation was conducted using eggNOG-mapper^89^ (v2.1.9, default parameters), based on eggNOG 5.0 orthology predictions and integrated databases including Pfam 33.1, KEGG, EC numbers, GO terms, and CAZy families^90^. Given the short length of smORFs, these eggNOG-based functional assignments should be interpreted as tentative annotations. Signal peptides were predicted with SignalP 6.0 (fast model)^91^, which identifies classical Sec/SPI signal peptides (SP), twin-arginine translocation (TAT) signals, lipoprotein signal peptides (LIPO and TATLIPO), and PILIN (Sec/SPIII), a specialized class of Sec pathway signal peptides. Transmembrane helices were predicted with TMHMM^92^ (v2.0, default parameters), allowing the determination of their number and topology.

### AMP prediction and physicochemical feature analysis

AMPs were predicted by ampir (https://ampir.marine-omics.net/; precursor peptide model), Macrel^93^ (v 1.5.0, default parameters), amPEPpy^94^ (v 1.1.0, parameter: --seed 2012), and AMPLify^95^ (v 2.0.0, parameter: -m imbalanced). Any peptide predicted as an AMP by at least one method was retained, and the combined outputs were merged to generate a comprehensive c_AMP catalog for each habitat. The amino acid composition of deep-sea c_AMPs was compared with experimentally validated AMPs from DRAMP^53^ (v4) and APD3 ^54^, as well as predicted datasets including AMPSphere^20^, EPs from the human proteome^55^, and AMPs identified from marine biofilms^21^. Additional physicochemical features were computed using custom Python scripts (Figshare: https://doi.org/10.6084/m9.figshare.30692342), including peptide length, molecular weight, isoelectric point (pI), net charge at pH 7 (NetCharge_pH7), GRAVY (grand average of hydropathy), amphiphilicity index, hydrophobic moment, linear moment, polyproline II (PPII) propensity, and disorder propensity. For visualization, 2,000 AMPs were randomly selected from each habitat-specific catalog and each reference database.

### Protein structure and peptide-protein interaction prediction

Three-dimensional structures of 131 selected c_AMPs were predicted using AlphaFold^96^ (v2.0; parameter: full_dbs), and model confidence was evaluated using pLDDT scores. Peptide–protein complex structures (targets: tRNA modifying enzymes TruA and TtcA) were generated using AlphaFold3 ^97^, and prediction quality were assessed using ipTM and pTM scores. All structural models were visualized and exported as images using PyMOL (http://www.pymol.org).

### Assessment of deep-sea smORF and c_AMP novelty

Each smORF catalog was compared against established small-protein reference databases, including sORFdb^35^, SmProt 2 ^36^, and GMSC^4^, to evaluate the homology and novelty of the identified deep-sea smORFs. Deep-sea c_AMPs were additionally compared with experimentally validated AMPs from DRAMP^53^ and APD3 ^54^, as well as predicted AMP datasets including AMPSphere^20^, EPs^55^, and marine biofilm AMPs ^21^. All pairwise comparisons were performed using DIAMOND^88^ (v2.1.8, parameters: --more-sensitive --evalue 1e-5).

### Deep-sea c_AMP selection for synthesis and testing

A total of 131 c_AMPs from the five deep-sea habitats were synthesized by solid-phase peptide synthesis using the Fmoc strategy by Sangon Biotech (Shanghai) Co., Ltd. Peptide molecular weights were verified by mass spectrometry, and purity (> 90%) was confirmed by high-performance liquid chromatography. The selection criteria for these peptides are described below.

Deep-sea c_AMPs were initially filtered by peptide length, excluding sequences longer than 40 amino acids, as conventional Fmoc-based solid-phase peptide synthesis often yields lower efficiencies and requires repeated recoupling steps for longer peptides^98, 99^. Two complementary groups of c_AMPs were then selected for downstream synthesis and functional testing: (i) peptides predicted to have high antimicrobial potential, and (ii) peptides randomly selected to represent taxonomic diversity across habitats. For the first group, we employed EvoGradient^100^ to predict and virtually evolve candidate bioactive peptides, an explainable deep learning framework integrating convolutional neural network (CNN)^101^, long short-term memory networks (LSTM)^102^, attention networks^103^ and Transformer models^103^. Peptides with high predicted antimicrobial activity were subsequently filtered based on solubility (> 0.5) using the criteria implemented in Protein–Sol^104^. This resulted in the selection of five peptides from abyssal plains, five from seamounts (one overlapping with the abyssal plain group), 16 from cold seeps, 18 from hydrothermal vents, and 18 from hadal trenches (one overlapping with the abyssal plain group). For the second group, c_AMPs were selected according to their taxonomic classification at a defined rank to ensure representation across taxonomic levels, retaining at least one peptide per rank. This yielded five c_AMPs from abyssal plains, four from seamounts, 21 from cold seeps, 21 from hydrothermal vents, and 20 from hadal trenches (one overlapping with the abyssal plain group).

### Bacterial and fungal inhibition experiment

Nine representative strains were selected for initial antimicrobial screening, including Gram-positive bacteria *P. filamentosa* and *S. aureus* (ATCC RN450); Gram-negative bacteria *E. coli* strains (DH5*α* and BW25113), *K. pneumoniae* (ATCC 13883), *V. parahaemolyticus* (VP.1997; ATCC 17802), *H. bluephagenesis* (TD01), and *A. pittii*; and the yeast *S. cerevisiae* (BY4741). Strains were streaked onto Luria–Bertani (LB) agar medium and incubated overnight at 37°C, except *H. bluephagenesis,* which was cultured on LB agar supplemented with 6 × NaCl (60 g L⁻¹ NaCl); *S. aureus*, which was grown on Mueller–Hinton (MH) agar; and *S. cerevisiae*, which was grown on yeast extract–peptone–dextrose (YPD) agar at 30°C. Single colonies were inoculated into appropriate liquid media and cultured overnight at 220 r.p.m. (37°C for bacteria; 30°C for *S. cerevisiae*). Overnight bacterial cultures were diluted 1:100 (v/v) into fresh medium and grown for 3 h to mid-exponential phase; *S. cerevisiae* cultures were diluted to an OD_600_ of 0.1 and grown for 6 h to reach mid-exponential phase. Bacterial suspensions were adjusted to an OD_600_ of 0.0002, and *S. cerevisiae* suspensions were adjusted to an OD_600_ of 0.02 (≈ 2 × 10^5^ CFU mL^−1^ in YPD or yeast extract–peptone– ethanol–glycerol (YPEG) medium) and used as inocula for testing 131 synthesized c_AMPs.

Freeze-dried c_AMPs was dissolved in double-distilled water to 0.4 mmol L^−1^. Three experimental groups were established in 96-well plates: (1) blank control, 100 μL fresh medium; (2) microbial control, 50 μL fresh medium and 50 μL bacterial or yeast suspension; and (3) c_AMPs treatment, 50 µL microbial suspension and 50 µL c_AMP solution (final concentration 125 µmol L^−1^). Each well contained a final volume of 100 µL and a microbial density of 1 × 10^5^ CFU mL⁻¹. Plates were incubated at 37°C for bacteria (30°C for *S. cerevisiae*) with shaking at 220 r.p.m for 16 h. OD_600_ values were normalized by (OD_600, experiment_ − OD_600, blank_) / (OD_600, control_ − OD_600, blank_). The c_AMPs were considered effective if they reduced bacterial or yeast growth by > 75% relative to controls. All assays were performed with three biological replicates, each including three technical replicates.

### Minimum inhibitory concentration determination

MICs of c_AMPs were determined by the broth microdilution method^19^. Briefly, microbial inocula were prepared as described for the inhibition experiment. A twofold serial dilution of each c_AMP (160–2.5 µM) was prepared in fresh medium, and 50 µL of each dilution was mixed with 50 μL of microbial inoculum in 96-well plates, resulting in final peptide concentrations ranging from 80 to 1.25 µM and a microbial density of 1 × 10^5^ CFU mL^−1^. Wells containing medium only (blank control) and microbial cultures without peptide (growth control) were included on each plate. The MIC was defined as the lowest peptide concentration with no detectable growth after 16 h incubation (37°C for bacteria; 30°C for *S. cerevisiae*). All assays were performed in triplicate with three independent biological replicates.

### Minimum bactericidal concentration determination

Minimum bactericidal concentrations (MBCs) of bioactive c_AMPs were determined according to the European Committee on Antimicrobial Susceptibility Testing (EUCAST) guidelines for bactericidal activity assessment. Briefly, bacterial cultures were prepared as described for MIC determination. After 16 h exposure to c_AMPs, 50 µL of culture from each well was spread onto LB agar plates lacking antimicrobial agents and incubated at 37°C for 18–24 h. *H. bluephagenesis* was plated onto LB agar supplemented with 6 × NaCl (60 g L^−1^ NaCl), *S. aureus* onto MH agar, and *S. cerevisiae* onto YPD or YPEG agar at 30°C. CFUs were enumerated, and the bactericidal rate was calculated as: Bactericidal rate (%) = (1 – CFU_sample_ / CFU_control_) × 100. The MBC was defined as the lowest peptide concentration resulting in ≥ 99.9% reduction in CFUs compared to the untreated control (corresponding to ≤ 0.1% survival). All assays were performed in triplicate.

### Cytotoxicity determination

Cytotoxicity was evaluated using the Cell Counting Kit-8 (C6005; NCM Biotech). HEK293T cells were cultured in Dulbecco’s Modified Eagle Medium (DMEM) supplemented with 10% fetal bovine serum (FBS) at 37°C in 5% CO_2_ for 24 h. Cells were then subcultured, diluted to 1 × 10^5^ cells mL^−1^, and 50 μL suspension was transferred to each well of a 96-well microplate. After 24 h, DMEM containing different concentrations of c_AMPs was dispensed to each well. Cells were treated with c_AMPs for 48 h. Subsequently, 10 µL of CCK-8 solution was added to each well and incubated for 30 min at 37°C, after which absorbance at 450 nm was measured using a microplate reader. Relative cell viability was calculated as (OD_450 sample_ – OD_450 blank_) / (OD_450 control_ – OD_450 blank_). All assays were performed with three biological replicates. The same procedure was applied to HepG2 and HeLa cells.

### Transmission electron microscopy examination

*P. filamentosa* cells were treated with three deep-sea c_AMPs (cold seep-32, cold seep-37, and hydrothermal vent-19) at a final concentration of 1 × MIC. In contrast, the control group received phosphate-buffered saline (PBS) under the same conditions. After incubation at 37°C for 2 h following previous studies^105^, bacterial suspensions were collected by centrifugation (6,000 r.p.m, 1 min, 4°C) and washed twice with PBS. Cell pellets were resuspended and immediately processed for ultrathin sectioning for observation. Samples were mounted on copper grids, negatively stained with 1% (w/v) uranyl acetate for 15 s, air-dried, and examined using a transmission electron microscope (TEM; Thermo Fisher Scientific Helios 5 UC) operated at 80 kV. Morphological alterations in cell walls and membranes were compared between peptide-treated and control cells.

### RNA sequencing and analysis

To investigate transcriptional responses induced by representative c_AMPs, *P. filamentosa* was treated with hydrothermal vent-19 (1 × MIC), *V. parahaemolyticus* with hydrothermal vent-19 (1 × MIC), *A. pittii* with cold seep-9 (1 × MIC), and *S. cerevisiae* with cold seep-38 (1 × MIC) for 4 h, each with three biological replicates. PBS was used as the control. Total RNA was extracted using TRIzol® reagent (Invitrogen) following the manufacturer’s instructions and treated with DNase I (TaKaRa) to remove residual genomic DNA. RNA integrity and purity were assessed using an Agilent 2100 Bioanalyzer (Agilent Technologies) and a NanoDrop ND-2000 spectrophotometer (Thermo Fisher Scientific). High-quality RNA samples (OD_260/280_ = 1.8–2.2, OD_260/230_ ≥ 2.0, RIN ≥ 6.5, 28S:18S ≥ 1.0, and total RNA ≥ 10 μg) were used for library construction.

For prokaryotic samples, rRNA was removed using the Ribo-Zero rRNA Removal Kit (Epicenter), and strand-specific libraries were constructed from 5 μg of total RNA following standard protocols. RNA was fragmented, reverse-transcribed into cDNA, end-repaired, adenylated, and ligated to indexed adaptors. Libraries were size-selected for 200–300 bp cDNA fragments using 2% Low Range Ultra Agarose and PCR-amplified with Phusion DNA polymerase (NEB). For the eukaryotic sample (*S. cerevisiae*), poly (A) + mRNA was purified using oligo (dT) magnetic beads, fragmented, and processed using the same workflow. Final libraries were quantified using a TBS380 fluorometer (Turner Biosystems) and sequenced on the MGI-T7 DNBSEQ platform (Shanghai BIOZERON Biotech) with 150 bp paired-end reads.

Raw reads were processed using Trimmomatic^106^ (v0.36; parameters: SLIDINGWINDOW:4:15 MINLEN:75) to remove adaptors, low-quality bases, and short reads. Reads containing more than 10% ambiguous bases (N) were discarded. Quality-controlled reads were further filtered against the rRNA database^107^ (Rfam) using sortmeRNA^108^ (v2.1) to remove rRNA contaminants. For bacterial samples, clean reads were aligned to the respective reference genomes using Rockhopper^109^. For *S. cerevisiae*, reads were aligned to the reference genome using HISAT2 ^110^, and gene-level counts were obtained using featureCounts (http://subread.sourceforge.net/). Gene expression levels were normalized as reads per kilobase of transcript per million mapped reads (RPKM)^111^. Differential expression and functional enrichment analyses were performed on the CFViSA^112^ platform. DEGs between treatment and control groups were identified using edgeR^113^ with thresholds of |log_2_(fold change)| ≥ 1 and false discovery rate (FDR) < 0.05. GO enrichment analysis was performed using Goatools^114^, and KEGG pathway enrichment was conducted using KOBAS^115^. GO terms^116^ and KEGGpathways^117^ with Bonferroni-corrected *P* < 0.05 were considered significantly enriched.

### Statistical analysis

Statistical analyses were performed using R v4.2.3. Data normality was assessed using the Shapiro–Wilk test prior to downstream analyses. The Kruskal–Wallis rank-sum test was used to compare physicochemical properties across deep-sea c_AMPs and reference AMP datasets. Pairwise comparisons were conducted using Wilcoxon rank-sum tests, and *P* values were adjusted using the Bonferroni correction. Corrected *P* < 0.05 was considered statistically significant.

### Data availability

All transcriptome sequencing data generated in this study have been deposited in Genome Sequence Archive (GSA) under accession number PRJCA052490. Sequence files for smORF and c_AMP datasets from the five deep-sea habitats have been deposited on Figshare (DOI: 10.6084/m9.figshare.30692342).

### Code availability

Original codes have been deposited at Figshare (DOI: 10.6084/m9.figshare.30692342). All computational tools used in this study were executed with default parameters unless otherwise specified, as detailed in the Methods.

## Acknowledgements

This work was supported by National Key R & D Program of China (2022YFC2805505, 2024YFA0916503); National Natural Science Foundation of China (42376115, 42406109, 32122050, 32370074); Natural Science Foundation of Fujian Province (2023J06042, 2025J011005); Scientific Research Foundation of Third Institute of Oceanography, MNR (2025013); and Hubei Hongshan Laboratory (CDSJ, 2025hsqd014). We thank Tianxueyu Zhang and Ruiqi Lin for assistance with data collection, Hai Zhang, Chen Ling, Xianming Deng for providing strains, and Qiang Zheng, Weipeng Zhang, Fabai Wu, and Rui Cheng for helpful discussions.

## Author contributions

X.D. and Z.L. conceived and designed the study. Q.J. and Y.H. carried out the omics analyses. Z.L. and C.Y. performed the experiments. Y.D., F.L., Z.H. and C.S. contributed to methodological development and discussions. Q.J., Y.H., X.D. and Z.L. wrote the manuscript with input from all authors. All authors reviewed and approved the final manuscript.

## References

1. Zhang, C., Peng, Y., Liu, X., Wang, J. & Dong, X. Deep-sea microbial genetic resources: new frontiers for bioprospecting. Trends in Microbiology 32, 321–324 (2024).

2. Wright, E.S. & Vetsigian, K.H. Inhibitory interactions promote frequent bistability among competing bacteria. Nature Communications 7, 11274 (2016).

3. Dong, X. et al. A vast repertoire of secondary metabolites potentially influences community dynamics and biogeochemical processes in cold seeps. Science Advances 10, eadl2281 (2024).

4. Duan, Y. et al. A catalog of small proteins from the global microbiome. Nature Communications 15, 7563 (2024).

5. Storz, G., Wolf, Y.I. & Ramamurthi, K.S. Small proteins can no longer be ignored. Annual Review of Biochemistry 83, 753–777 (2014).

6. Gaßel, M., Möllenkamp, T., Puppe, W. & Altendorf, K. The KdpF subunit is part of the K^+^-translocating Kdp complex of *Escherichia coli* and is responsible for stabilization of the complex *in vitro*. Journal of Biological Chemistry 274, 37901–37907 (1999).

7. Salazar, M.E., Podgornaia, A.I. & Laub, M.T. The small membrane protein MgrB regulates PhoQ bifunctionality to control PhoP target gene expression dynamics. Molecular Microbiology 102, 430–445 (2016).

8. Lloyd, C.R., Park, S., Fei, J. & Vanderpool, C.K. The small protein SgrT controls transport activity of the glucose-specific phosphotransferase system. Journal of Bacteriology 199, 10.1128/jb.00869-00816 (2017).

9. Alix, E. & Blanc-Potard, A.B. Hydrophobic peptides: novel regulators within bacterial membrane. Molecular Microbiology 72, 5–11 (2009).

10. Sassone-Corsi, M. et al. Microcins mediate competition among *Enterobacteriaceae* in the inflamed gut. Nature 540, 280–283 (2016).

11. Lazzaro, B.P., Zasloff, M. & Rolff, J. Antimicrobial peptides: Application informed by evolution. Science 368, eaau5480 (2020).

12. Yu, G., Baeder, D.Y., Regoes, R.R. & Rolff, J. Predicting drug resistance evolution: insights from antimicrobial peptides and antibiotics. Proceedings of the Royal Society B: Biological Sciences 285, 20172687 (2018).

13. Kintses, B. et al. Phylogenetic barriers to horizontal transfer of antimicrobial peptide resistance genes in the human gut microbiota. Nature Microbiology 4, 447–458 (2019).

14. Ahrens, C.H., Wade, J.T., Champion, M.M. & Langer, J.D. A practical guide to small protein discovery and characterization using mass spectrometry. Journal of Bacteriology 204, e00353–00321 (2022).

15. Hyatt, D. et al. Prodigal: prokaryotic gene recognition and translation initiation site identification. BMC Bioinformatics 11, 1–11 (2010).

16. Aspden, J.L. et al. Extensive translation of small open reading frames revealed by Poly-Ribo-Seq. eLife 3, e03528 (2014).

17. Leong, A.Z.-X. et al. Short open reading frames (sORFs) and microproteins: an update on their identification and validation measures. Journal of Biomedical Science 29, 19 (2022).

18. Rodrjguez del Rjo, Á., et al. Functional and evolutionary significance of unknown genes from uncultivated taxa. Nature 626, 377–384 (2024).

19. Ma, Y. et al. Identification of antimicrobial peptides from the human gut microbiome using deep learning. Nature Biotechnology 40, 921–931 (2022).

20. Santos-Júnior, C.D. et al. Discovery of antimicrobial peptides in the global microbiome with machine learning. Cell 187, 3761–3778.e3716 (2024).

21. Fan, S. et al. Bioprospecting of culturable marine biofilm bacteria for novel antimicrobial peptides. iMeta 3 (2024).

22. Han, Y. et al. A comprehensive genomic catalog from global cold seeps. Scientific Data 10, 596 (2023).

23. Zhang, C. et al. The majority of microorganisms in gas hydrate-bearing subseafloor sediments ferment macromolecules. Microbiome 11, 37 (2023).

24. Liang, X., Nong, X.-H., Huang, Z.-H. & Qi, S.-H. Antifungal and antiviral cyclic peptides from the deep-sea-derived fungus *Simplicillium obclavatum* EIODSF 020. Journal of Agricultural and Food Chemistry 65, 5114–5121 (2017).

25. Zhou, X. et al. Marthiapeptide A, an anti-infective and cytotoxic polythiazole cyclopeptide from a 60 L scale fermentation of the deep sea-derived *Marinactinospora thermotolerans* SCSIO 00652. Journal of Natural Products 75, 2251–2255 (2012).

26. Casella, V. et al. Novel insights into the *Nobilamide* family from a deep-sea *Bacillus*: Chemical diversity, biosynthesis and antimicrobial activity towards multidrug-resistant bacteria. Marine Drugs 23, 41 (2025).

27. Lai, C. et al. New polyketides from a hydrothermal vent sediment fungus Trichoderma sp. JWM29-10-1 and their antimicrobial effects. Marine Drugs 20, 720 (2022).

28. Wei, X., Hu, Y., Sun, C. & Wu, S. Characterization of a novel antimicrobial peptide bacipeptin against foodborne pathogens. Journal of Agricultural and Food Chemistry 72, 5283–5292 (2024).

29. Hancock, R.E.W. & Lehrer, R. Cationic peptides: a new source of antibiotics. Trends in Biotechnology 16, 82–88 (1998).

30. Koehbach, J. & Craik, D.J. The vast structural diversity of antimicrobial peptides. Trends in Pharmacological Sciences 40, 517–528 (2019).

31. Oliveira Junior, N.G., Souza, C.M., Buccini, D.F., Cardoso, M.H. & Franco, O.L. Antimicrobial peptides: structure, functions and translational applications. Nature Reviews Microbiology 23, 687–700 (2025).

32. Sneideris, T. et al. Targeting nucleic acid phase transitions as a mechanism of action for antimicrobial peptides. Nature Communications 14, 7170 (2023).

33. Seefeldt, A.C. et al. Structure of the mammalian antimicrobial peptide Bac7(1–16) bound within the exit tunnel of a bacterial ribosome. Nucleic Acids Research 44, 2429–2438 (2016).

34. Ghosh, A. et al. Indolicidin targets duplex DNA: structural and mechanistic insight through a combination of spectroscopy and microscopy. ChemMedChem 9, 2052–2058 (2014).

35. Hahnfeld, J.M. et al. sORFdb–a database for sORFs, small proteins, and small protein families in bacteria. BMC Genomics 26, 110 (2025).

36. Li, Y. et al. SmProt: A reliable repository with comprehensive annotation of small proteins identified from ribosome profiling. *Genomics*, Proteomics & Bioinformatics 19, 602–610 (2021).

37. Xiao, X. et al. Microbial ecosystems and ecological driving forces in the deepest ocean sediments. Cell 188, 1363–1377.e1369 (2025).

38. Dick, G.J. The microbiomes of deep-sea hydrothermal vents: distributed globally, shaped locally. Nature Reviews Microbiology 17, 271–283 (2019).

39. Orcutt, B.N., Sylvan, J.B., Knab, N.J. & Edwards, K.J. Microbial ecology of the dark ocean above, at, and below the seafloor. Microbiol Mol Biol Rev 75, 361–422 (2011).

40. Nunoura, T. et al. Microbial diversity in sediments from the bottom of the Challenger Deep, the Mariana Trench. Microbes and Environments 33, 186–194 (2018).

41. Schauberger, C. et al. Microbial community structure in hadal sediments: high similarity along trench axes and strong changes along redox gradients. The ISME Journal 15, 3455–3467 (2021).

42. Jiao, Y., Yang, S. & Bao, W. Biogeographic patterns and community assembly mechanisms of bacterial community in the upper seawater of seamounts and non-seamounts in the Eastern Indian Ocean. Applied and Environmental Microbiology 90, e01424–01424 (2024).

43. Li, H., Zhou, H., Yang, S. & Dai, X. Stochastic and deterministic assembly processes in seamount microbial communities. Applied and Environmental Microbiology 89, e00701–00723 (2023).

44. Mirdita, M., Steinegger, M., Breitwieser, F., Söding, J. & Levy Karin, E. Fast and sensitive taxonomic assignment to metagenomic contigs. Bioinformatics 37, 3029–3031 (2021).

45. Parks, D.H. et al. GTDB: an ongoing census of bacterial and archaeal diversity through a phylogenetically consistent, rank normalized and complete genome-based taxonomy. Nucleic Acids Research 50, D785–D794 (2022).

46. Li, Y., Cao, W., Wang, Y. & Ma, Q. Microbial diversity in the sediments of the southern Mariana Trench. Journal of Oceanology and Limnology 37, 1024–1029 (2019).

47. Tully, B.J., Graham, E.D. & Heidelberg, J.F. The reconstruction of 2,631 draft metagenome-assembled genomes from the global oceans. Scientific Data 5, 170203 (2018).

48. Zhou, Y.-L., Mara, P., Cui, G.-J., Edgcomb, V.P. & Wang, Y. Microbiomes in the Challenger Deep slope and bottom-axis sediments. Nature Communications 13, 1515 (2022).

49. Jiang, Q. et al. Cold seeps are potential hotspots of deep-sea nitrogen loss driven by microorganisms across 21 phyla. Nature Communications 16, 1646 (2025).

50. Galperin, M.Y. et al. COG database update: focus on microbial diversity, model organisms, and widespread pathogens. Nucleic Acids Research 49, D274–D281 (2021).

51. Miravet-Verde, S., et al. Unraveling the hidden universe of small proteins in bacterial genomes. Molecular Systems Biology 15, e8290 (2019).

52. Yadavalli, S.S. & Yuan, J. Bacterial small membrane proteins: the swiss army knife of egulators at the lipid bilayer. Journal of Bacteriology 204, e00344–00321 (2022).

53. Ma, T. et al. DRAMP 4.0: an open-access data repository dedicated to the clinical translation of antimicrobial peptides. Nucleic Acids Research 53, D403–D410 (2024).

54. Wang, G., Li, X. & Wang, Z. APD3: the antimicrobial peptide database as a tool for research and education. Nucleic Acids Research 44, D1087–D1093 (2015).

55. Torres, M.D.T. et al. Mining for encrypted peptide antibiotics in the human proteome. Nature Biomedical Engineering 6, 67–75 (2022).

56. Le, C.-F., Fang, C.-M. & Sekaran, S.D. Intracellular targeting mechanisms by antimicrobial peptides. Antimicrobial Agents and Chemotherapy 61, 10.1128/aac.02340-02316 (2017).

57. Zhang, Q.-Y. et al. Antimicrobial peptides: mechanism of action, activity and clinical potential.Military Medical Research 8, 48 (2021).

58. Melino, S. et al. Metal-binding and nuclease activity of an antimicrobial peptide analogue of the salivary histatin 5. Biochemistry 45, 15373–15383 (2006).

59. Migliolo, L. et al. Structural and functional characterization of a multifunctional alanine-rich peptide analogue from *Pleuronectes americanus*. PLOS ONE 7, e47047 (2012).

60. Yacoub, H.A., Al-Maghrabi, O.A., Ahmed, E.S. & Uversky, V.N. Abundance and functional roles of intrinsic disorder in the antimicrobial peptides of the NK-lysin family. Journal of Biomolecular Structure and Dynamics 35, 836–856 (2017).

61. Balleza, D. The role of flexibility in the bioactivity of short *α*-helical antimicrobial peptides. ntibiotics 14, 422 (2025).

62. Tuerkova, A. et al. Effect of helical kink in antimicrobial peptides on membrane pore formation. eLife 9, e47946 (2020).

63. Cotter, P.D., Ross, R.P. & Hill, C. Bacteriocins—a viable alternative to antibiotics? Nature Reviews Microbiology 11, 95–105 (2013).

64. Reddy, K.V.R., Yedery, R.D. & Aranha, C. Antimicrobial peptides: premises and promises. International Journal of Antimicrobial Agents 24, 536–547 (2004).

65. Bass, D. et al. Yeast forms dominate fungal diversity in the deep oceans. Proceedings of the Royal Society B: Biological Sciences 274, 3069–3077 (2007).

66. Yang, S., Xu, W., Gao, Y., Chen, X. & Luo, Z.-H. Fungal diversity in deep-sea sediments from Magellan seamounts environment of the western Pacific revealed by high-throughput Illumina sequencing. Journal of Microbiology 58, 841–852 (2020).

67. Gaser, O.A. et al. Alteration of metabolic activity regulates mitochondrial temperature in diagnosis in HepG2 hepatocellular carcinoma cells. Scientific Reports 15, 41155 (2025).

68. Hancock, R.E.W. & Sahl, H.-G. Antimicrobial and host-defense peptides as new anti-infective therapeutic strategies. Nature Biotechnology 24, 1551–1557 (2006).

69. Zhong, C. et al. An antimicrobial peptide as a potential therapy for bacterial pneumonia that alleviates antimicrobial resistance. Nature Communications 16, 10488 (2025).

70. Giacomucci, S., Cros, C.D.-N., Perron, X., Mathieu-Denoncourt, A. & Duperthuy, M. Flagella-dependent inhibition of biofilm formation by sub-inhibitory concentration of polymyxin B in Vibrio cholerae. PLOS ONE 14, e0221431 (2019).

71. Plaza, N., Pérez-Reytor, D., Corsini, G., García, K. & Urrutia, Í.M. Contribution of the type III secretion system (T3SS2) of *Vibrio parahaemolyticus* in mitochondrial stress in human intestinal cells. Microorganisms 12, 813 (2024).

72. Antonietti, M. et al. An analysis of intrinsic protein disorder in antimicrobial peptides. The Protein Journal 44, 175–191 (2025).

73. Busch, D.J. et al. Intrinsically disordered proteins drive membrane curvature. Nature Communications 6, 7875 (2015).

74. Gong, H. et al. How do antimicrobial peptides disrupt the lipopolysaccharide membrane leaflet of Gram-negative bacteria? Journal of Colloid and Interface Science 637, 182–192 (2023).

75. Li, J. et al. Deep sea cold seeps are a sink for mercury and source for methylmercury. Communications Earth & Environment 5, 324 (2024).

76. Dong, X. et al. Metabolic potential of uncultured bacteria and archaea associated with petroleum seepage in deep-sea sediments. Nature Communications 10, 1816 (2019).

77. Dong, X. et al. Phylogenetically and catabolically diverse diazotrophs reside in deep-sea cold seep sediments. Nature Communications 13, 4885 (2022).

78. Dong, X. et al. Thermogenic hydrocarbon biodegradation by diverse depth-stratified microbial populations at a Scotian Basin cold seep. Nature Communications 11, 5825 (2020).

79. Jiang, Q., Jing, H., Jiang, Q. & Zhang, Y. Insights into carbon-fixation pathways through metagonomics in the sediments of deep-sea cold seeps. Marine Pollution Bulletin 176, 113458 (2022).

80. Lu, R. et al. Asgard archaea in the haima cold seep: Spatial distribution and genomic insights. Deep Sea Research Part I: Oceanographic Research Papers 170, 103489 (2021).

81. Li, W.-L., Wu, Y.-Z., Zhou, G.-w., Huang, H. & Wang, Y. Metabolic diversification of anaerobic methanotrophic archaea in a deep-sea cold seep. Marine Life Science & Technology 2, 431–441 (2020).

82. Xiao, X. et al. Metal-driven anaerobic oxidation of methane as an important methane sink in methanic cold seep sediments. Microbiology Spectrum 11, e05337–05322 (2023).

83. Chen, S., Zhou, Y., Chen, Y. & Gu, J. fastp: an ultra-fast all-in-one FASTQ preprocessor. Bioinformatics 34, i884–i890 (2018).

84. Li, D., Liu, C.-M., Luo, R., Sadakane, K. & Lam, T.-W. MEGAHIT: an ultra-fast single-node solution for large and complex metagenomics assembly via succinct de Bruijn graph. Bioinformatics 31, 1674–1676 (2015).

85. Durrant, M.G. & Bhatt, A.S. Automated prediction and annotation of small open reading frames in microbial genomes. Cell Host & Microbe 29, 121–131.e124 (2021).

86. Fu, L., Niu, B., Zhu, Z., Wu, S. & Li, W. CD-HIT: accelerated for clustering the next-generation sequencing data. Bioinformatics 28, 3150–3152 (2012).

87. Steinegger, M. & Söding, J. MMseqs2 enables sensitive protein sequence searching for the analysis of massive data sets. Nature Biotechnology 35, 1026–1028 (2017).

88. Buchfink, B., Reuter, K. & Drost, H.G. Sensitive protein alignments at tree-of-life scale using DIAMOND. Nature Methods 18, 366–368 (2021).

89. Cantalapiedra, C.P., Hernández-Plaza, A., Letunic, I., Bork, P. & Huerta-Cepas, J. eggNOG-mapper v2: functional annotation, orthology assignments, and domain prediction at the metagenomic scale. Molecular Biology and Evolution 38, 5825–5829 (2021).

90. Huerta-Cepas, J. et al. eggNOG 5.0: a hierarchical, functionally and phylogenetically annotated orthology resource based on 5090 organisms and 2502 viruses. Nucleic Acids Research 47, D309–D314 (2019).

91. Teufel, F. et al. SignalP 6.0 predicts all five types of signal peptides using protein language models. Nature Biotechnology 40, 1023–1025 (2022).

92. Krogh, A., Larsson, B., von Heijne, G. & Sonnhammer, E.L.L. Predicting transmembrane protein topology with a hidden markov model: application to complete genomes11Edited by F. Cohen. Journal of Molecular Biology 305, 567–580 (2001).

93. Santos-Junior, C.D., Pan, S., Zhao, X.-M. & Coelho, L.P. Macrel: antimicrobial peptide screening in genomes and metagenomes. PeerJ 8, e10555 (2020).

94. Lawrence, T.J. et al. amPEPpy 1.0: a portable and accurate antimicrobial peptide prediction tool. Bioinformatics 37, 2058–2060 (2020).

95. Li, C. et al. AMPlify: attentive deep learning model for discovery of novel antimicrobial peptides effective against WHO priority pathogens. BMC Genomics 23, 77 (2022).

96. Jumper, J. et al. Highly accurate protein structure prediction with AlphaFold. Nature 596, 583–589 (2021).

97. Abramson, J. et al. Accurate structure prediction of biomolecular interactions with AlphaFold 3. Nature 630, 493–500 (2024).

98. Kochendoerfer, G.G. & Kent, S.B.H. Chemical protein synthesis. Current Opinion in Chemical Biology 3, 665–671 (1999).

99. Palomo, J.M. Solid-phase peptide synthesis: an overview focused on the preparation of biologically relevant peptides. RSC Advances 4, 32658–32672 (2014).

100. Wang, B. et al. Explainable deep learning and virtual evolution identifies antimicrobial peptides with activity against multidrug-resistant human pathogens. Nature Microbiology 10, 332–347 (2025).

101. O’Shea, K. & Nash, R. An introduction to convolutional neural networks. ArXiv Preprint at https://arxiv.org/abs/1511.08458 (2015).

102. Hochreiter, S. & Schmidhuber, J. Long short-term memory. Neural Computation 9, 1735–1780 (1997).

103. Vaswani, A. et al. in Proceedings of the 31st International Conference on Neural Information Processing Systems 6000–6010 (Curran Associates Inc., Long Beach, California, USA; 2017).

104. Hebditch, M., Carballo-Amador, M.A., Charonis, S., Curtis, R. & Warwicker, J. Protein–Sol: a web tool for predicting protein solubility from sequence. Bioinformatics 33, 3098–3100 (2017).

105. Wang, T. et al. The effect of structural modification of antimicrobial peptides on their antimicrobial activity, hemolytic activity, and plasma stability. Journal of Peptide Science 27, e3306 (2021).

106. Bolger, A.M., Lohse, M. & Usadel, B. Trimmomatic: a flexible trimmer for Illumina sequence data. Bioinformatics 30, 2114–2120 (2014).

107. Sellqs Vidal, L., Ayala, R., Stan, G.-B. & Ledesma-Amaro, R. rfaRm: An R client-side interface to facilitate the analysis of the Rfam database of RNA families. PLOS ONE 16, e0245280 (2021).

108. Kopylova, E., Noé, L. & Touzet, H. SortMeRNA: fast and accurate filtering of ribosomal RNAs in metatranscriptomic data. Bioinformatics 28, 3211–3217 (2012).

109. Slezacek, J., Quillfeldt, P., Kaiya, H., Hykollari, A. & Fusani, L. Circulating profile of the appetite-regulating hormone ghrelin during moult-fast and chick provisioning in southern rockhopper penguins (*Eudyptes chrysocome chrysocome*). Hormones and Behavior 164, 105592 (2024).

110. Thakur, V. Transcriptome Data Analysis. (Springer US, New York, NY; 2024).

111. Ke, Y. et al. The progressive application of single-cell RNA sequencing technology in cardiovascular diseases. Biomedicine & Pharmacotherapy 154, 113604 (2022).

112. Wang, N. et al. CFViSA: A comprehensive and free platform for visualization and statistics in omics-data. Computers in Biology and Medicine 171, 108206 (2024).

113. Sarker, B., Matiur Rahaman, M., Alamin, M.H., Ariful Islam, M. & Nurul Haque Mollah, M. Boosting edgeR (Robust) by dealing with missing observations and gene-specific outliers in RNA-Seq profiles and its application to explore biomarker genes for diagnosis and therapies of ovarian cancer. Genomics 116, 110834 (2024).

114. Klopfenstein, D.V. et al. GOATOOLS: A python library for gene ontology analyses. Scientific Reports 8, 10872 (2018).

115. Bu, D. et al. KOBAS-i: intelligent prioritization and exploratory visualization of biological functions for gene enrichment analysis. Nucleic Acids Research 49, W317–W325 (2021).

116. Yan, H., Wang, S., Liu, H., Mamitsuka, H. & Zhu, S. GORetriever: reranking protein-description-based GO candidates by literature-driven deep information retrieval for protein function annotation. Bioinformatics 40, ii53-ii61 (2024).

117. Yaning, L. et al. Cpd861 targeting BCL2 to alleviate hepatic fibrosis: Network pharmacology, mendelian randomization, and molecular docking mechanisms. Current Pharmaceutical Design 30, 3291–3310 (2024).

