## Supplementary Figures 1-19 for "A global deep-sea small protein atlas reveals a reservoir of noncanonical antimicrobial peptides"

Supplementary Table legends

**Supplementary Table 1.** Sampling and sequencing metadata for all metagenomic datasets used in this study.

**Supplementary Table 2.** Comprehensive information for all smORF catalogs.

**Supplementary Table 3.** Summary statistics of smORFs shared between each pair of samples in five deep-sea habitats.

**Supplementary Table 4.** Detailed information on smORFs shared across all five deep-sea habitats. The table presents pairwise comparison data between the cold seep and each of the other four habitats.

**Supplementary Table 5.** COG functional distribution of shared small proteins across all five deep-sea habitats.

**Supplementary Table 6.** Relative abundance of archaeal and bacterial smORFs at the phylum level across different deep-sea habitats.

**Supplementary Table 7.** Relative abundance of archaeal and bacterial smORFs at the class level across different deep-sea habitats.

**Supplementary Table 8.** COG functional distribution of small proteins from the hadal trench, cold seep, hydrothermal vent, abyssal plain, and seamount.

**Supplementary Table 9.** Predicted transmembrane and secretory characteristics of small proteins across the five deep-sea habitats.

**Supplementary Table 10.** Amino acid frequency profiles of deep-sea c_AMPs compared with experimentally validated AMPs (databases, DRAMP and APD3) and computationally predicted AMPs (Eps and Biofilm).

**Supplementary Table 11.** Physicochemical properties of peptides from deep-sea c_AMPs compared with experimentally verified AMPs (databases, DRAMP and APD3) and computationally predicted AMPs (EPs and Biofilm).

**Supplementary Table 12.** Statistical analysis of pairwise comparisons of physicochemical properties among deep-sea c_AMPs and reference peptide datasets.

**Supplementary Table 13.** Relative OD_600_ values from bacterial inhibition assays.

**Supplementary Table 14.** Gradient-based determination of minimum inhibitory concentrations (MIC, μM).

**Supplementary Table 15.** Minimum inhibitory concentrations (MICs) of synthetic antimicrobial peptides against non-pathogenic and pathogenic microbial strains.

**Supplementary Table 16.** Minimum bactericidal concentrations (MBCs) of synthetic antimicrobial peptides against non-pathogenic and pathogenic microbial strains.

**Supplementary Table 17.** Cytotoxicity of synthetic antimicrobial peptides (c_AMPs) in mammalian cells, reported as normalized OD_450_ values.

**Supplementary Table 18.** Differential gene expression profile of *Acinetobacter pittii* in response to cold seep-9 treatment.

**Supplementary Table 19.** Differential gene expression profile of *Pseudomonas filamentosa* after exposure to hydrothermal vent-19.

**Supplementary Table 20.** Differential gene expression profile of *Vibrio parahaemolyticus* after treatment with hydrothermal vent-19.

**Supplementary Table 21.** Differential gene expression profile of *Saccharomyces cerevisiae* in response to cold seep-38 treatment.

Supplementary Figures

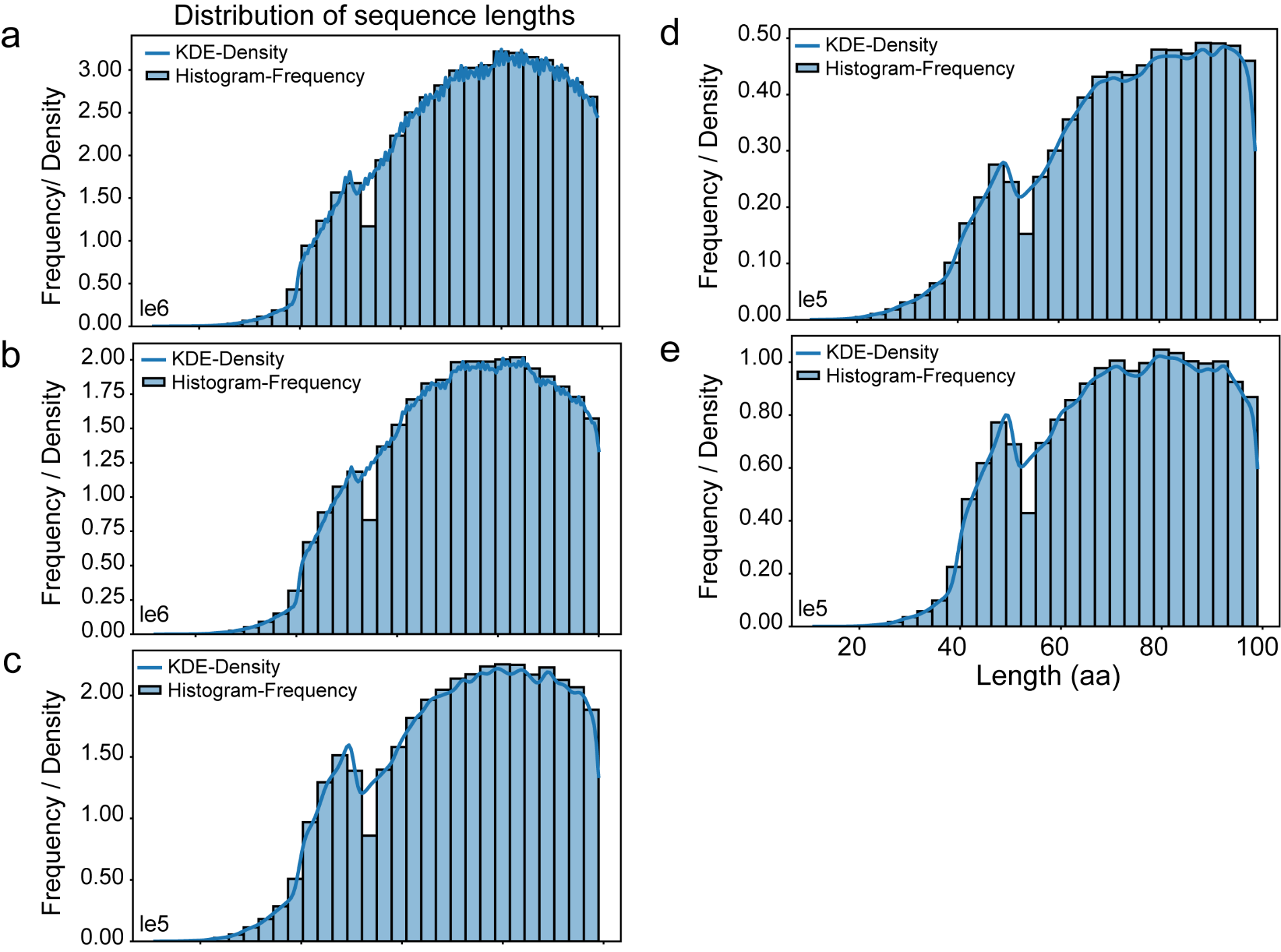

**Supplementary Figure 1.** **Summary statistics of deep-sea microbial smORF catalogs.** Distribution of smORF-encoded peptide lengths across five deep-sea habitats. Histograms overlaid with kernel density estimates illustrate the length profiles of smORFs identified from the hadal trench **(a)**, cold seep **(b)**, hydrothermal vent **(c)**, abyssal plain **(d)**, and seamount **(e)** datasets.

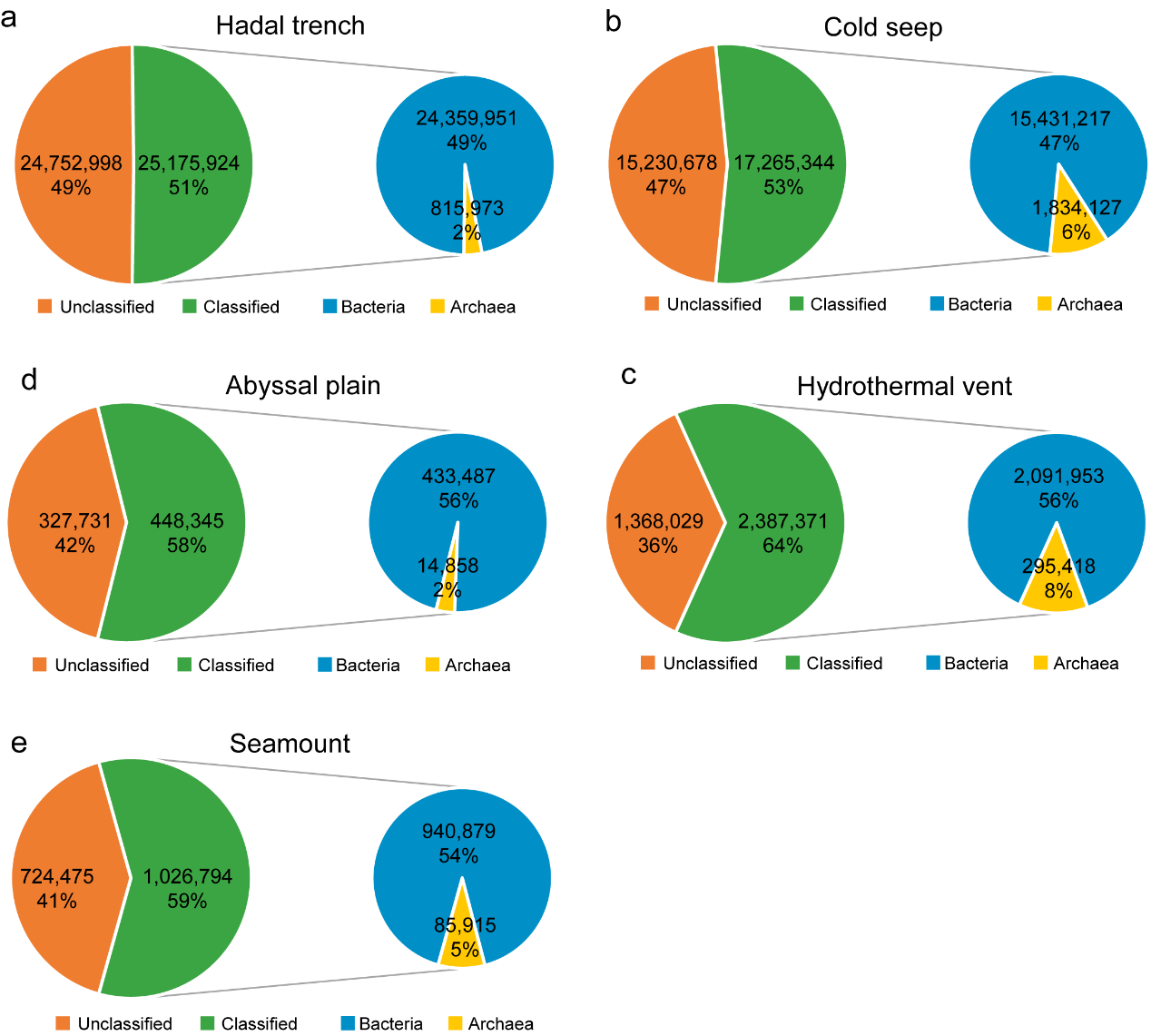

**Supplementary** **Figure 2.** **Taxonomic annotation of deep-sea microbial smORF catalogs.** Taxonomic assignments of smORFs were generated using the GTDB database (R220) for five deep-sea habitats: hadal trench **(a)**, cold seep **(b)**, hydrothermal vent **(c)**, abyssal plain **(d)**, and seamount **(e)**. For classified smORFs, the relative contributions of Bacteria and Archaea are shown in the magnified pie charts. Detailed taxonomic annotations for all smORFs across the five habitats are provided in **Supplementary Table 2**.

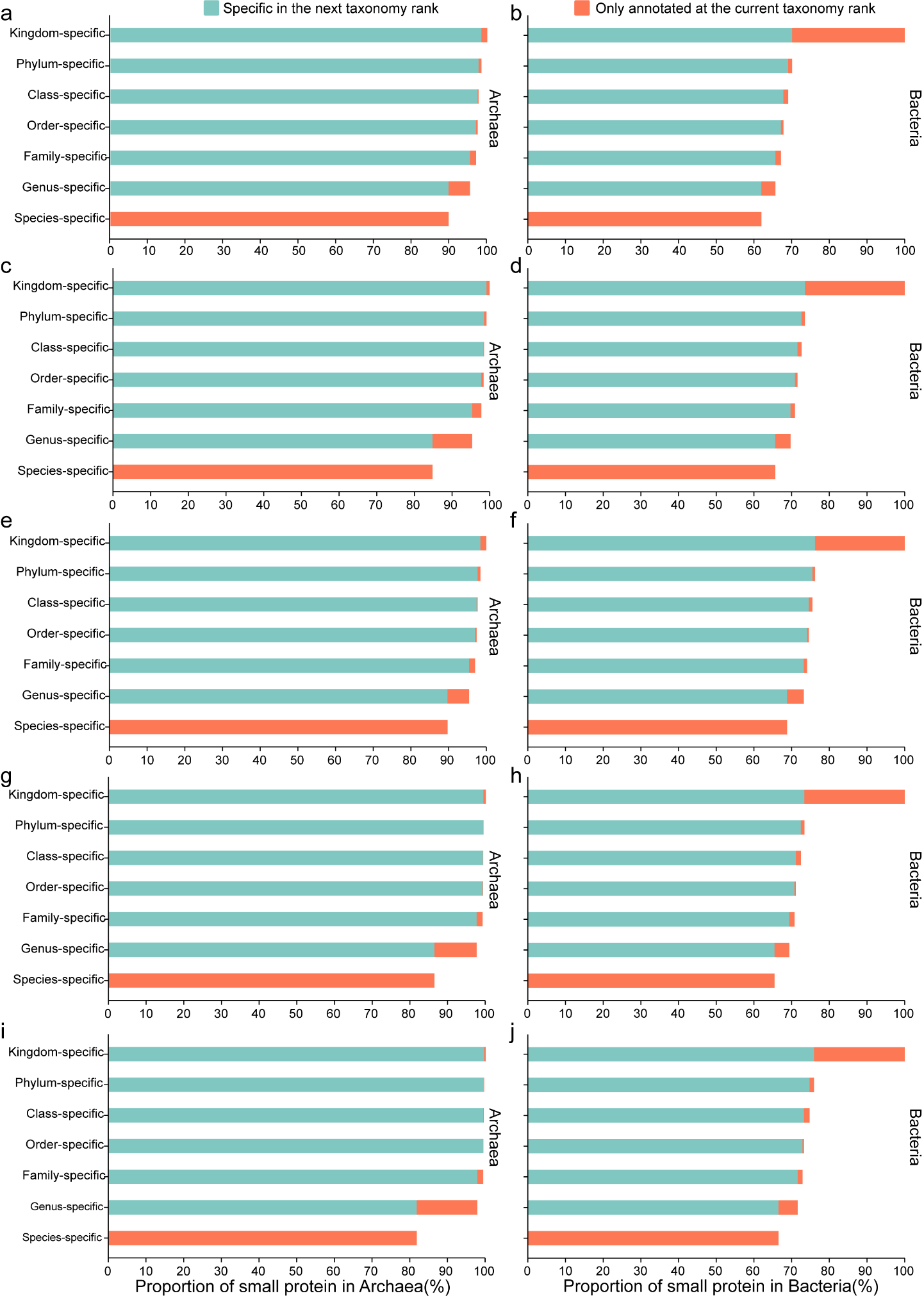

**Supplementary** **Figure 3.** **Taxonomic annotation resolution of smORFs across hierarchical ranks.** For each deep-sea habitat, including hadal trench **(a–b)**, cold seep **(c–d)**, hydrothermal vent **(e–f)**, abyssal plain **(g–h)**, and seamount **(i–j)**, the taxonomic resolution of smORFs was assessed across hierarchical ranks from kingdom to species. At a given rank, an smORF was considered assigned to that rank if all taxonomically annotated hits placed it within the same lineage. We further distinguish two cases, namely whether its members are annotated to the next taxonomic rank (light blue, marked specific in the next taxonomic rank), or not annotated to next rank (orange, marked only annotated at the current taxonomic rank). Other ranks are treated analogously (until we reach the level of species).

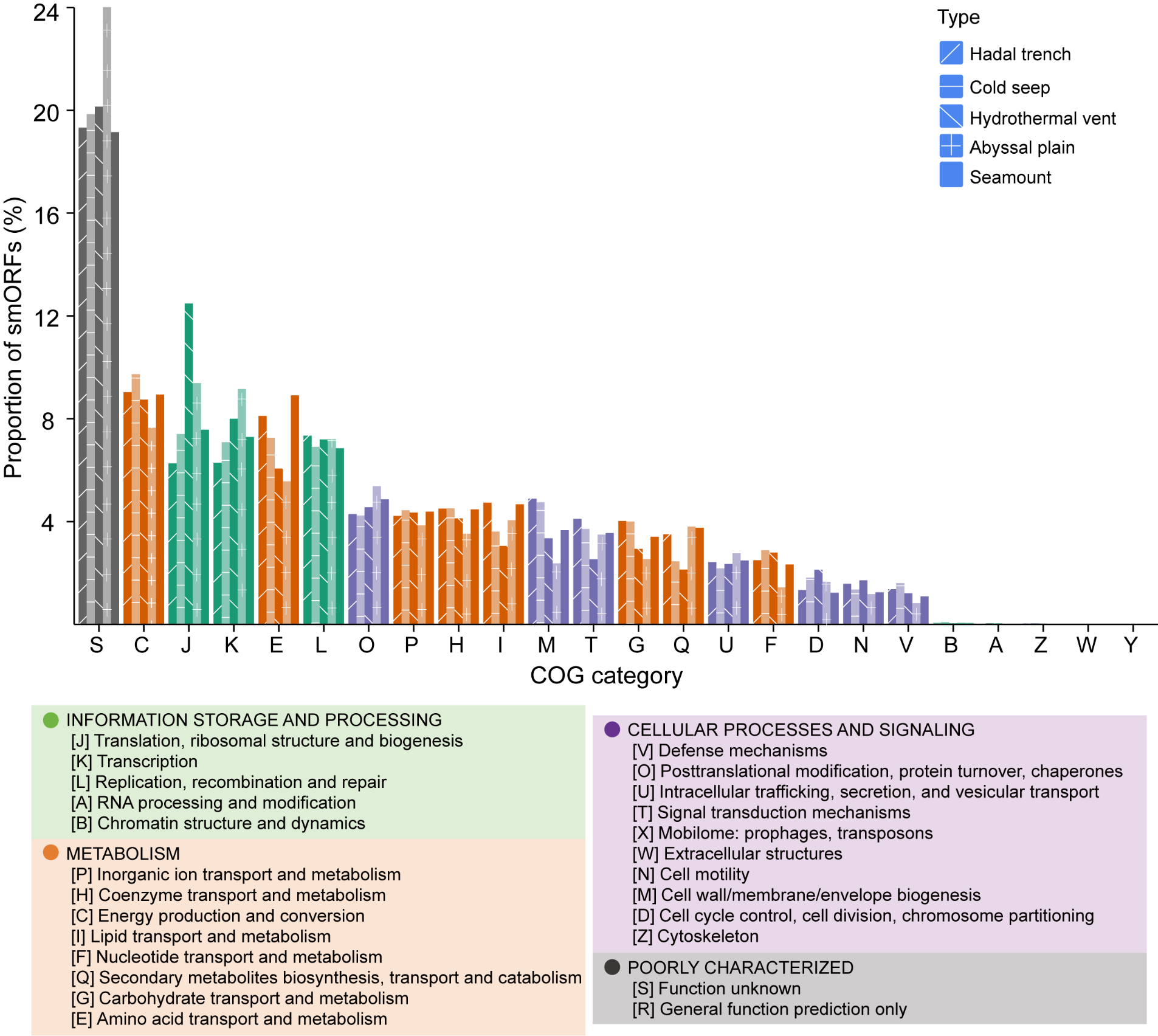

**Supplementary Figure 4. Functional annotation and structural features of smORF-encoded small proteins across deep-sea habitats.** COG-based functional classifications of smORF-encoded small proteins from five deep-sea habitats (hadal trench, cold seep, hydrothermal vent, abyssal plain, and seamount). Bars represent the proportional distribution of COG categories for archaeal and bacterial small proteins within each habitat.

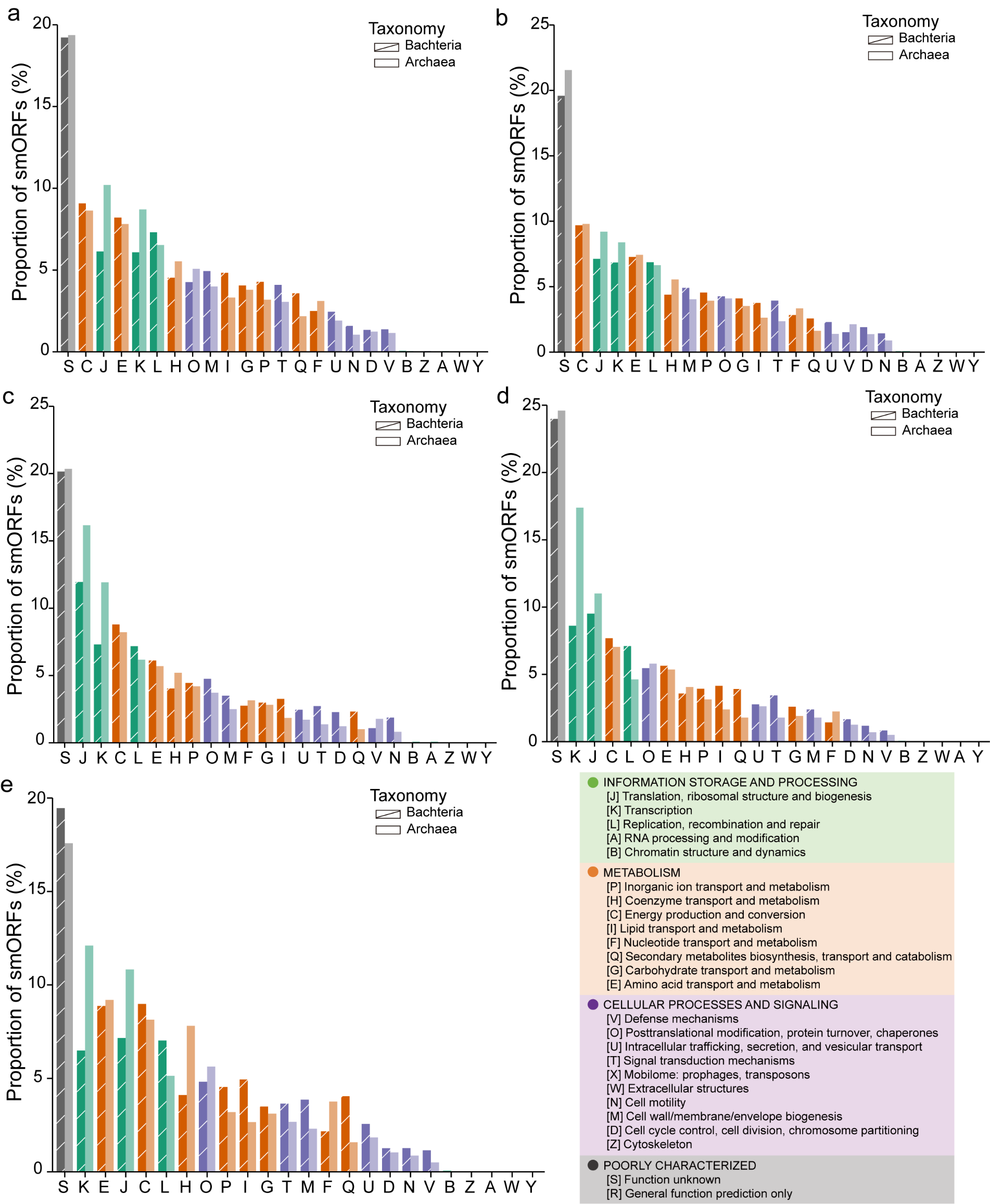

**Supplementary** **Figure 5.** **COG functional category distributions of archaeal and bacterial smORF-encoded small proteins across deep-sea habitats.** COG-based functional profiles of archaeal and bacterial small proteins are shown for five deep-sea habitats: hadal trench **(a)**, cold seep **(b)**, hydrothermal vent **(c)**, abyssal plain **(d)**, and seamount **(e)**. Bars represent the proportion of smORFs assigned to each COG category within the archaeal and bacterial lineages in each habitat.

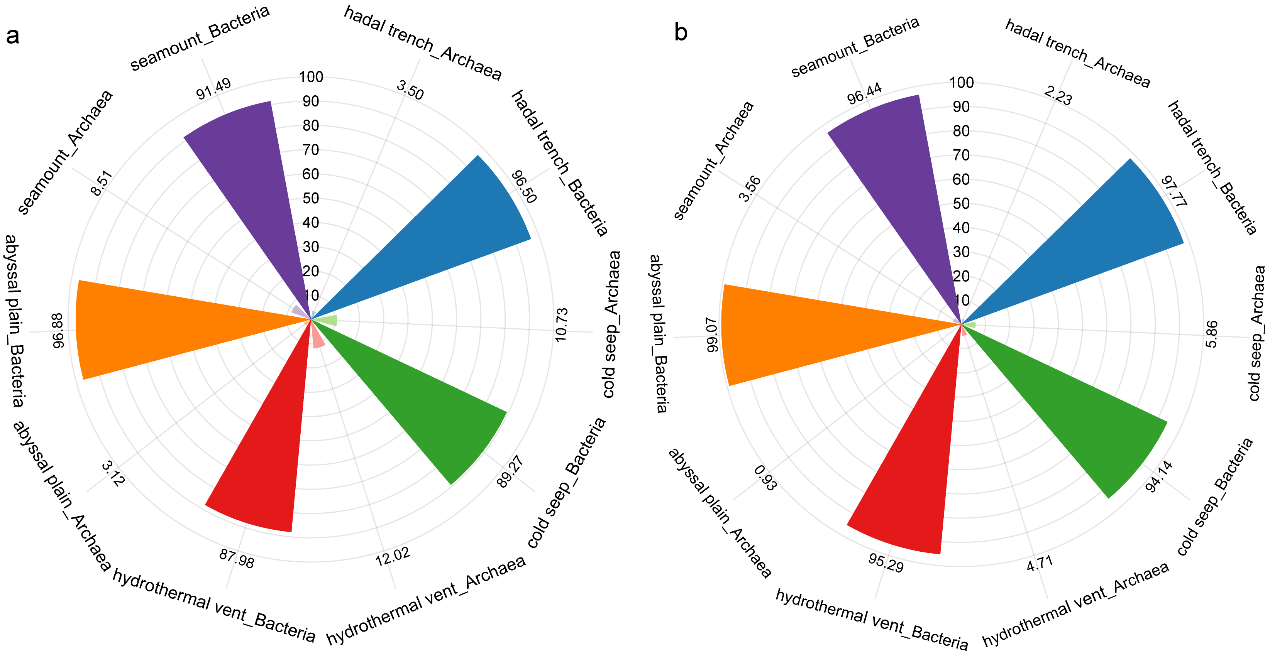

**Supplementary Figure 6. Transmembrane and signal-peptide features of archaeal and bacterial smORF-encoded small proteins across deep-sea habitats.** Radial plots showing the proportions of smORF-derived small proteins predicted to contain transmembrane domains **(a)** or signal peptides **(b)** within archaeal and bacterial lineages across five deep-sea habitats (hadal trench, cold seep, hydrothermal vent, abyssal plain, and seamount). Percentages represent the fraction of small proteins with the corresponding structural feature within each taxonomic group in each habitat.

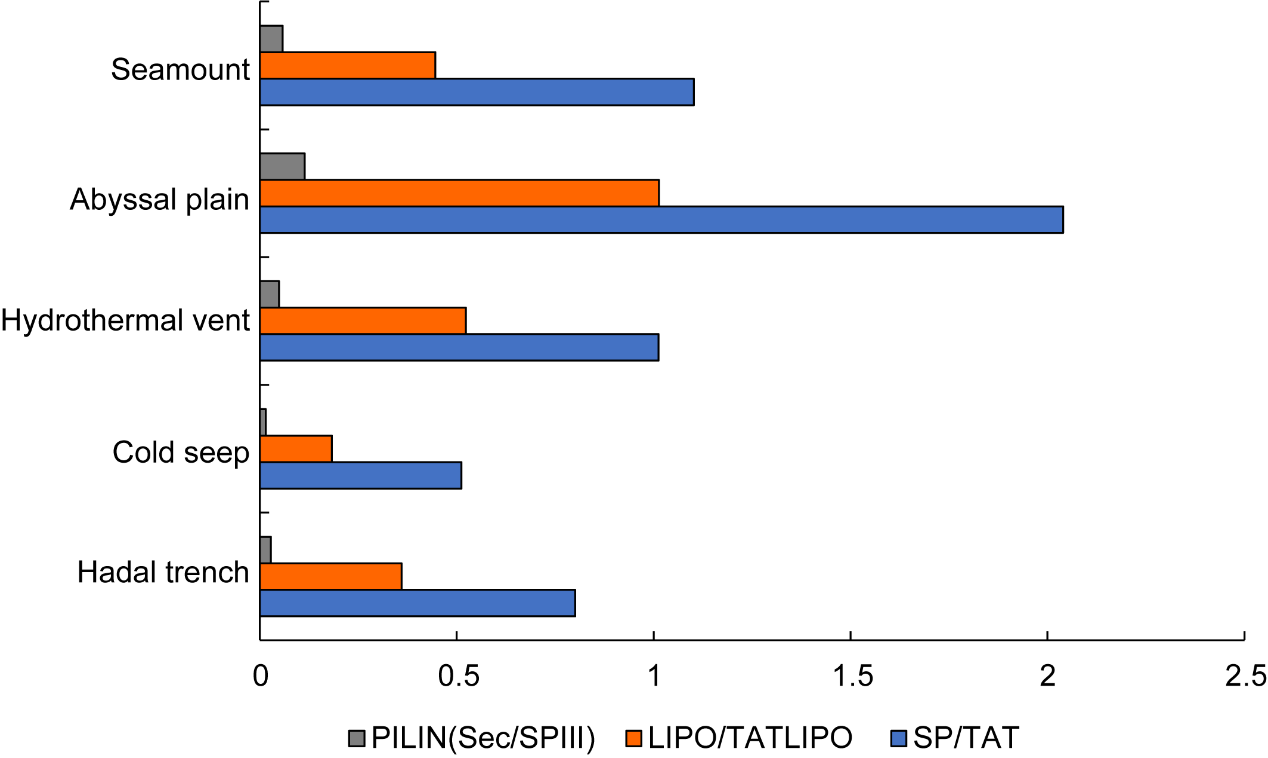

**Supplementary Figure 7. Distribution of signal peptide types in smORF-encoded small proteins across deep-sea habitats.** Proportions of smORF-derived small proteins predicted to contain distinct classes of signal peptides are shown for five deep-sea habitats. SP/TAT (blue): signal peptides associated with the Sec/SPI pathway or the twin-arginine translocation (TAT) pathway. LIPO/TATLIPO (orange): lipoprotein signal peptides processed by SPII, including TAT-dependent lipoprotein signal peptides. PILIN (Sec/SPIII) (grey): a specialized Sec pathway signal peptide processed by SPIII, typically found in type IV pilin and pilin-like proteins. Values represent the percentage of small proteins containing each signal peptide type within each habitat.

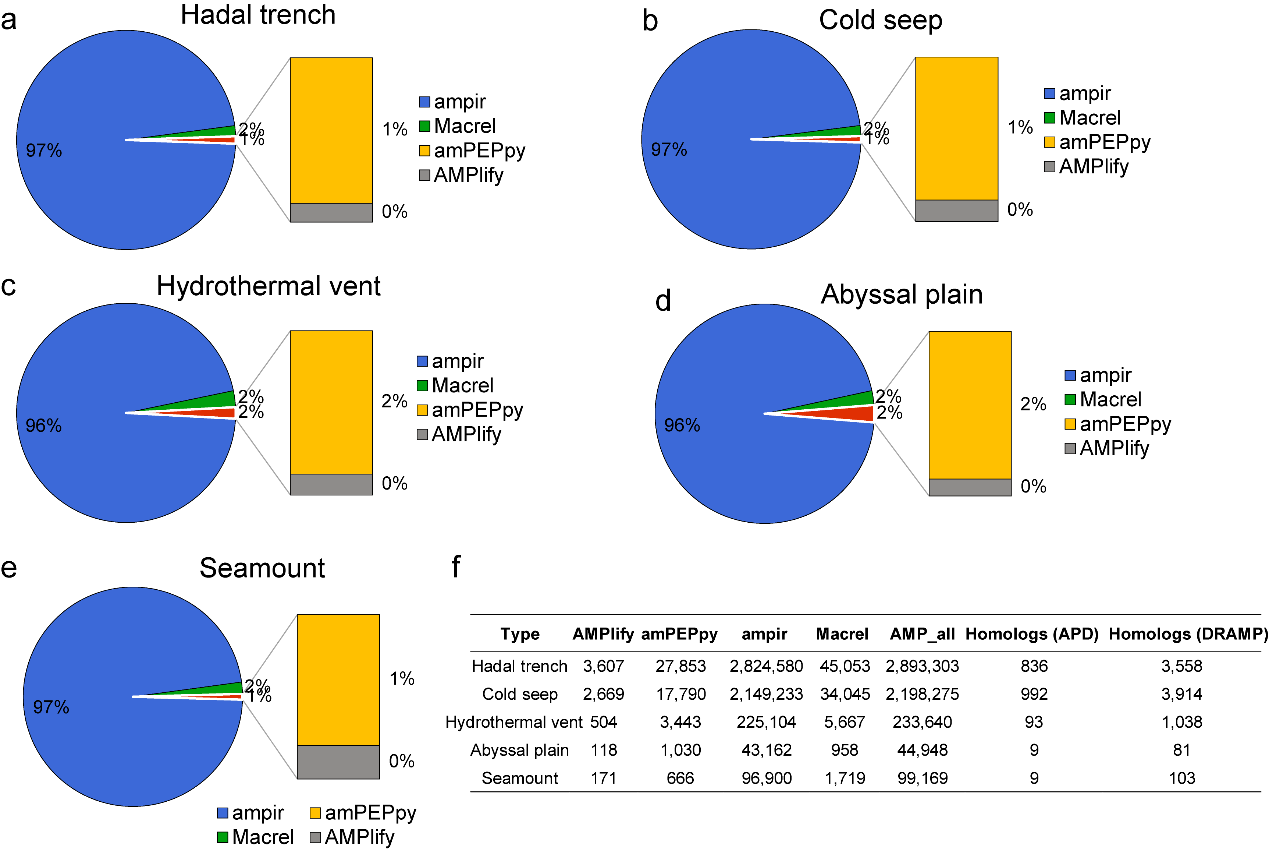

**Supplementary Figure 8. AMP candidate prediction across deep-sea habitats using four computational tools. (a–e)** Numbers and proportions of AMP candidates predicted by four software tools in the hadal trench (a), cold seep (b), hydrothermal vent (c), abyssal plain (d), and seamount (e) habitats. Insets illustrate the relative contribution of each tool to the combined AMP catalog within each habitat. **(f)** Summary of AMP predictions and homology assessments relative to experimentally validated AMPs in APD3 and DRAMP 4.0. AMP_all represents the non-redundant union of AMP candidates predicted by the four tools. The table reports the number of AMP candidates detected by each method and the number of homologs identified in each database. Because some peptides are predicted by multiple tools, the total number of AMP candidates is not equal to the sum of tool-specific predictions.

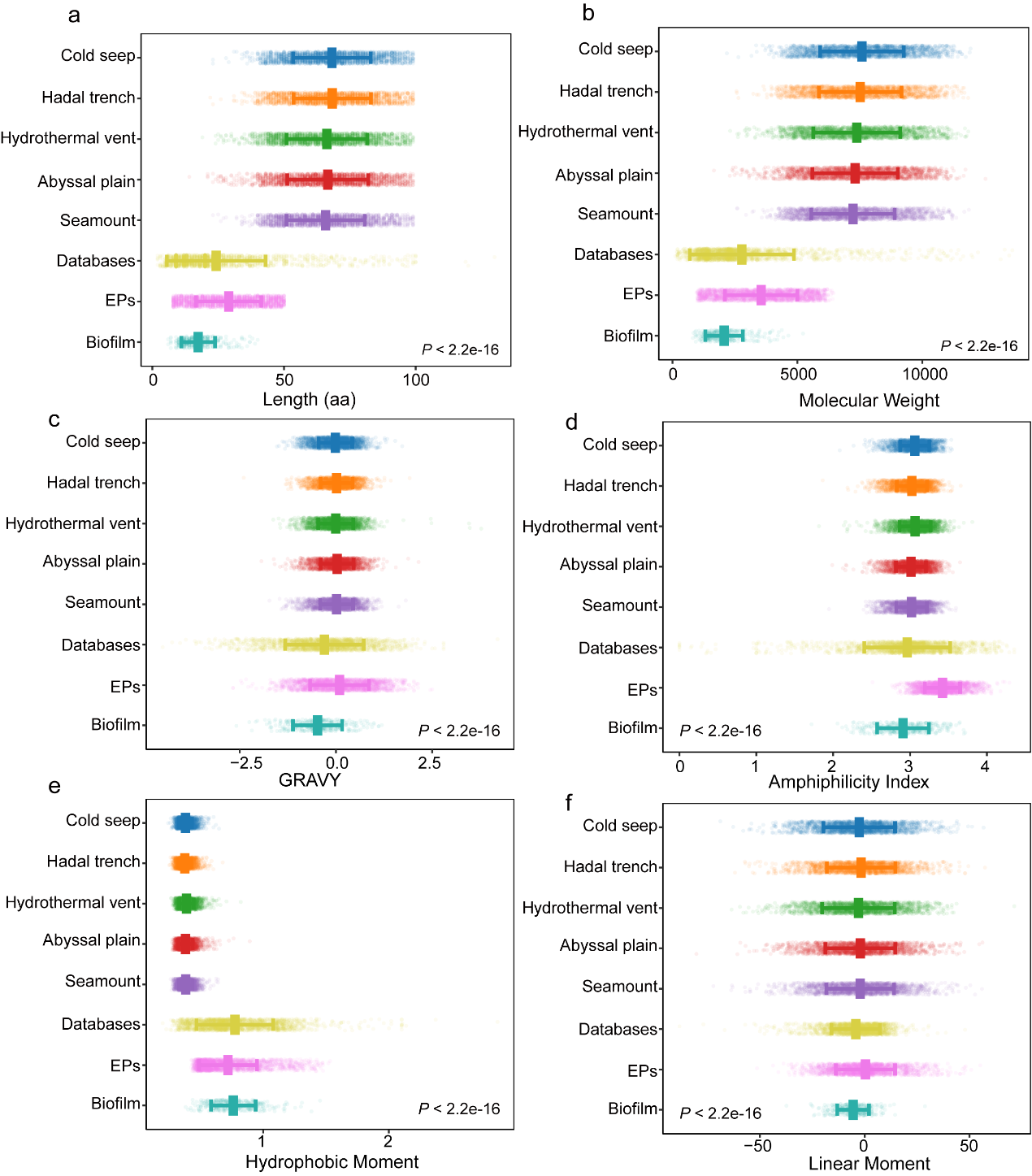

**Supplementary Figure 9. Distinct physicochemical properties of deep-sea smORF-derived antimicrobial peptides compared with reference peptide datasets. (a-e)** Multidimensional comparison of six physicochemical features across deep-sea c_AMPs and reference peptide datasets (databases (DRAMP + APD3), EPs, marine biofilm AMPs). These features inlucde peptide length (a), molecular weight (b), GRAVY (hydrophobicity; c), amphiphilicity index (d), hydrophobic moment (e), and linear moment (f). Deep-sea c_AMPs exhibit significantly distinct physicochemical profiles (*P* < 2.2 × 10^-16^), occupying a unique region of antimicrobial peptide sequence space.

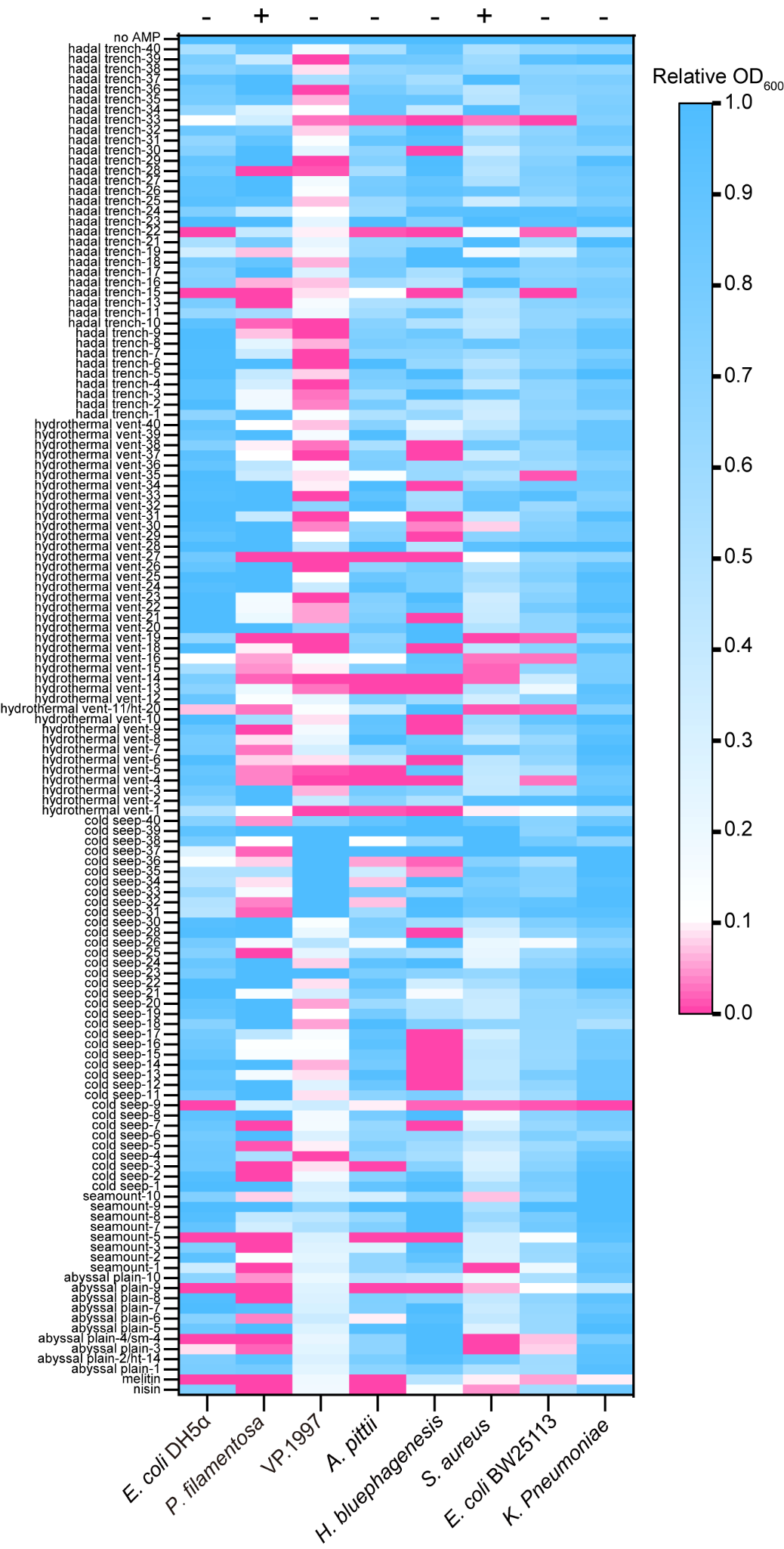

**Supplementary Figure 10. Inhibitory spectrum of 131 synthetic c_AMPs against microbial strains.** Heatmap showing the relative growth (OD_600_) of eight bacterial strains following treatment with 131 synthetic c_AMPs at 125 μM. Values represent OD_600_ normalized to untreated controls, with lower values (pink) indicating stronger growth inhibition. The tested strains include both Gram-positive (+) and Gram-negative (–) bacteria, as indicated at the top. Many peptides exhibit broad-spectrum inhibitory activity, with several achieving ≥75% reduction in OD_600_ relative to controls.

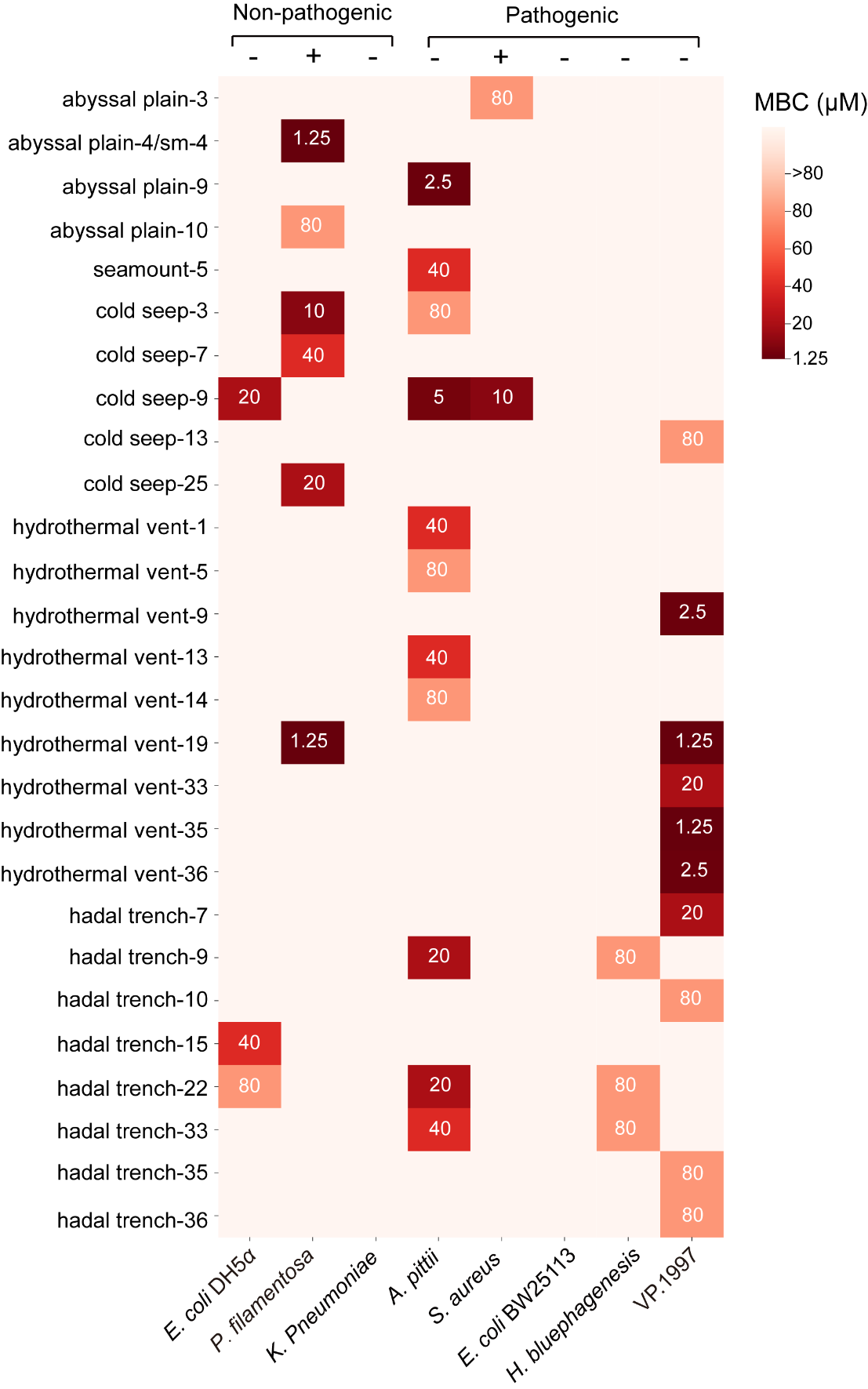

**Supplementary Figure 11. Minimum bactericidal concentrations (MBCs) of synthetic deep-sea c_AMPs against microbial strains.** Heatmap showing the MBC values (μM) of selected synthetic c_AMPs tested against eight bacterial strains, including both non-pathogenic and pathogenic species. Gram-positive (+) and Gram-negative (–) bacteria are indicated at the top. MBC values represent the lowest peptide concentration resulting in ≥99.9% reduction in viable cells relative to untreated controls. Assays were conducted in three independent biological replicates.

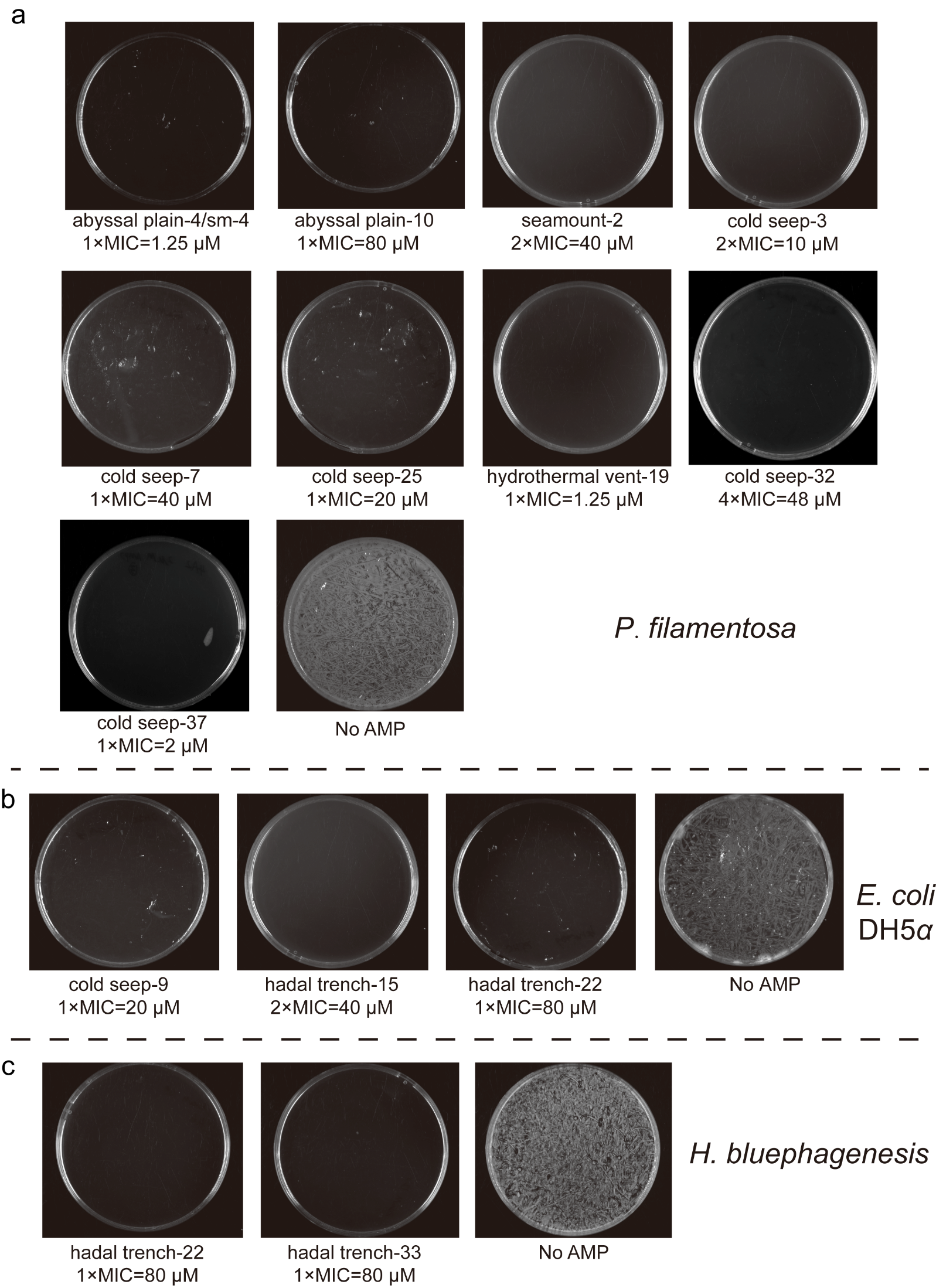

**Supplementary Figure 12. Representative plate images from minimum bactericidal concentration (MBC) assays of synthetic deep-sea c_AMPs.** Representative agar plate images showing bactericidal activity of selected synthetic c_AMPs against **(a)** *Priestia filamentosa*, **(b)** *Escherichia coli* DH5*α*, and **(c)** *Halomonas bluephagenesis*. For each peptide, cultures were plated after 16 h exposure to the indicated peptide concentration (expressed as multiples of the MIC). “No AMP” denotes untreated controls exhibiting full bacterial growth. All assays were performed in three independent biological replicates.

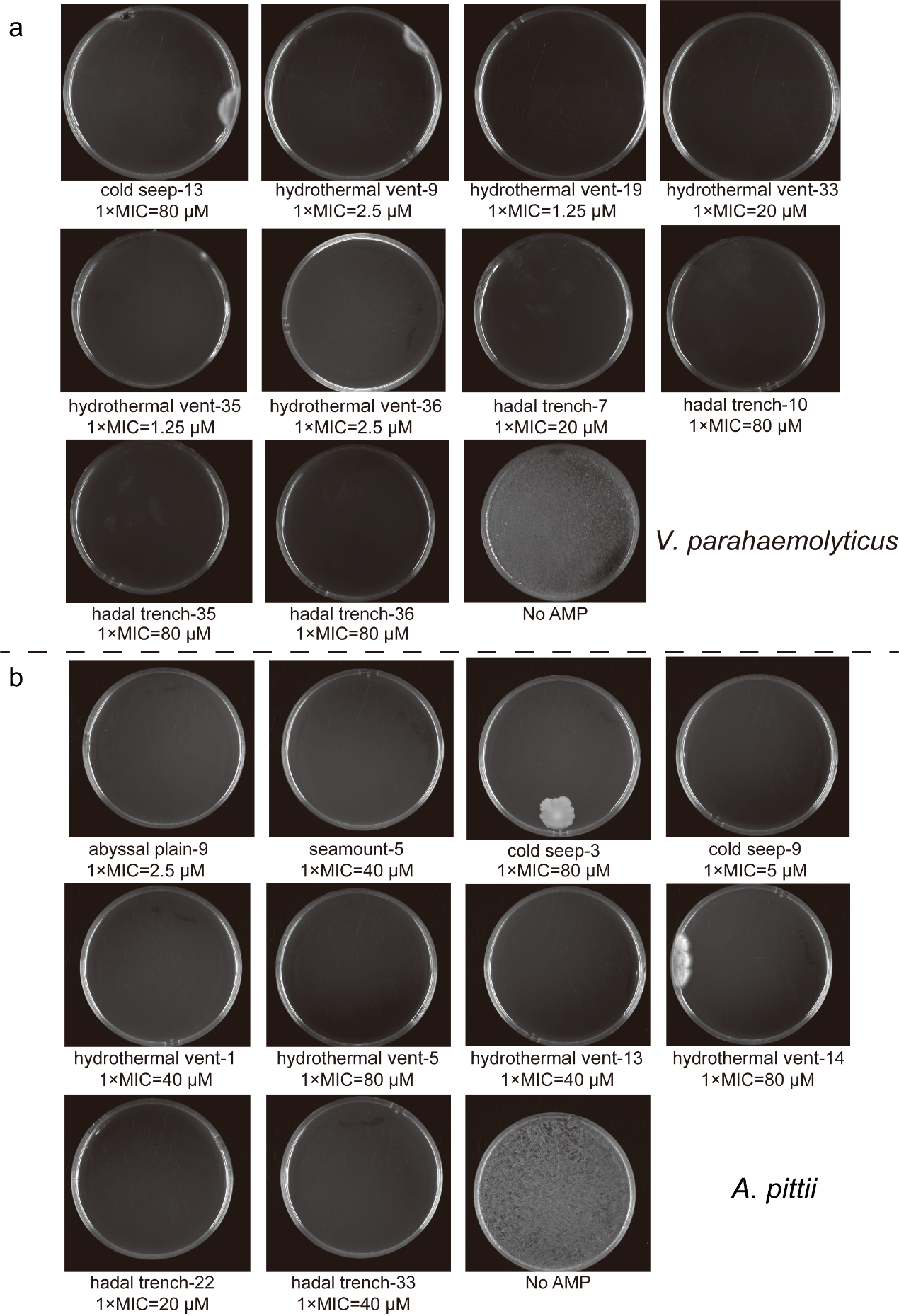

**Supplementary Figure 13. Representative plate images from minimum bactericidal concentration (MBC) assays of synthetic deep-sea c_AMPs.** Representative agar plate images showing bactericidal activity of selected synthetic c_AMPs against **(a)** *Vibrio parahaemolyticus* and **(b)** *Acinetobacter pittii*. For each peptide, cultures were plated after 16 h exposure to the indicated peptide concentration (expressed as multiples of the MIC). “No AMP” denotes untreated controls displaying full bacterial growth. All assays were performed in three independent biological replicates.

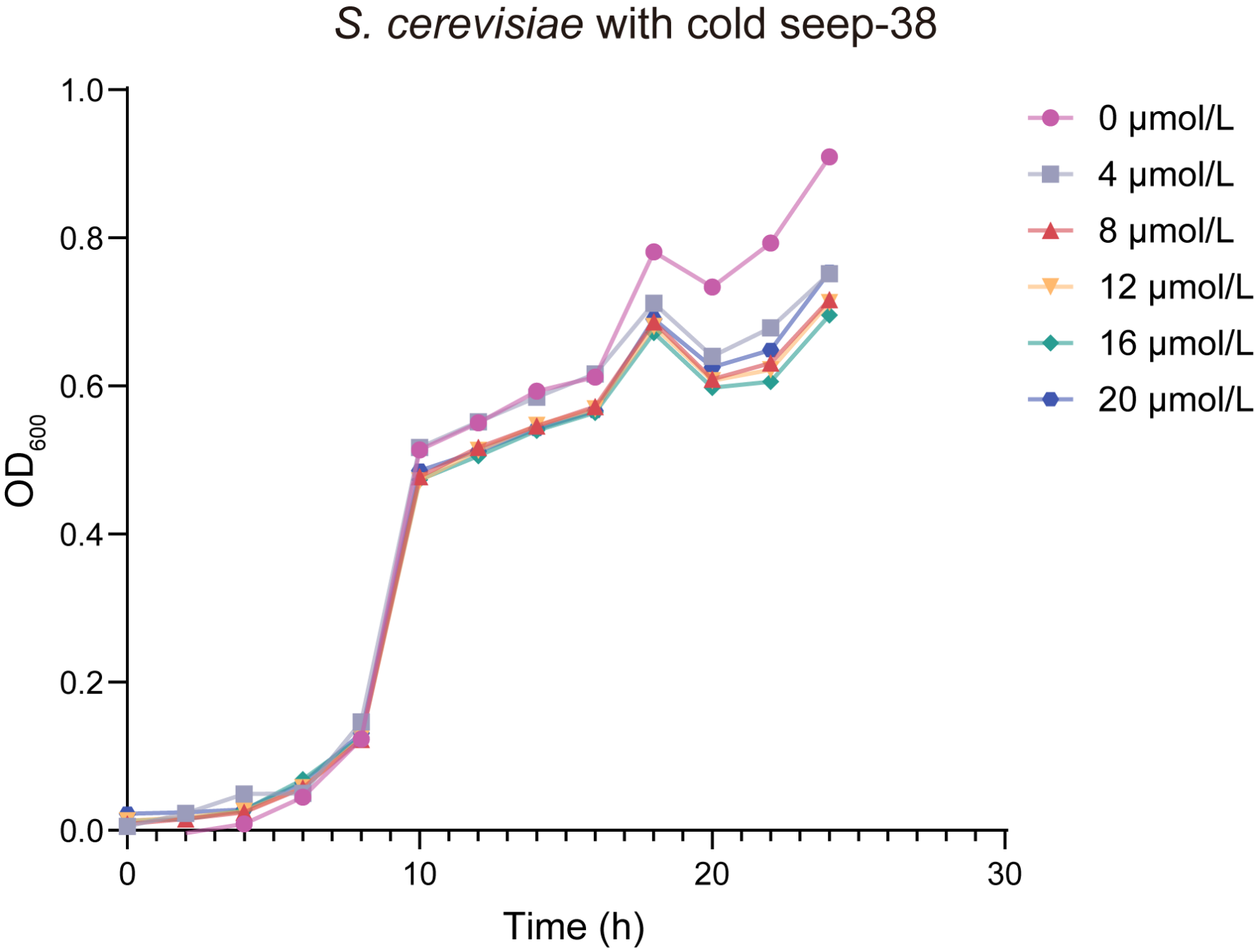

**Supplementary Figure 14. Growth curves of *S. cerevisiae* exposed to increasing concentrations of peptide cold seep-38.** Growth curves (OD_600_) of *S. cerevisiae* cultured in YPD medium in the presence of increasing concentrations of peptide cold seep-38. The control (0 μmol L⁻¹) is shown in pink, with treatment concentrations of 4 μmol L⁻¹ (grey), 8 μmol L⁻¹ (red), 12 μmol L⁻¹ (orange), 16 μmol L⁻¹ (green), and 20 μmol L⁻¹ (dark blue). The x-axis indicates incubation time (h).

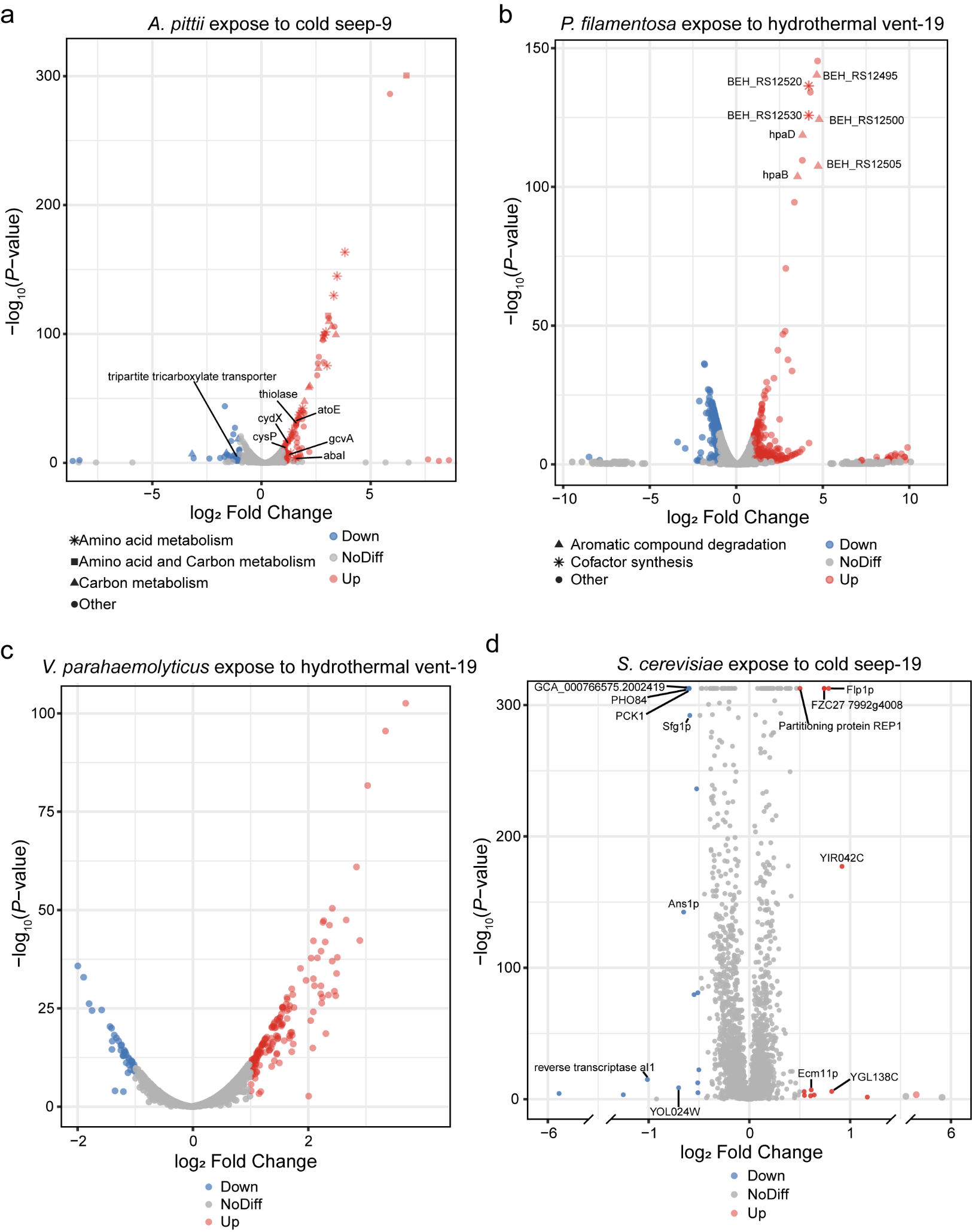

**Supplementary Figure 15. Volcano plots showing differential gene expression in microbial strains treated with deep-sea–derived antimicrobial peptides (c_AMPs). (a)** *Acinetobacter pittii* treated with peptide cold seep-9. **(b)** *Priestia filamentosa* treated with peptide hydrothermal vent-19. **(c)** *Vibrio parahaemolyticus* treated with peptide hydrothermal vent-19. **(d)** *Saccharomyces cerevisiae* treated with peptide cold seep-19, with selected genes labelled for clarity. Volcano plots display log_2_ fold change (x-axis) versus − log_10_(*P* value) (y-axis) for genes differentially expressed following AMP treatment. Significantly upregulated genes are shown in red, significantly downregulated genes in blue, and non-differentially expressed genes in grey (*P* < 0.05; thresholds of |log_2_FC| > 1 for panels a–c and > 0.5 for panel d).

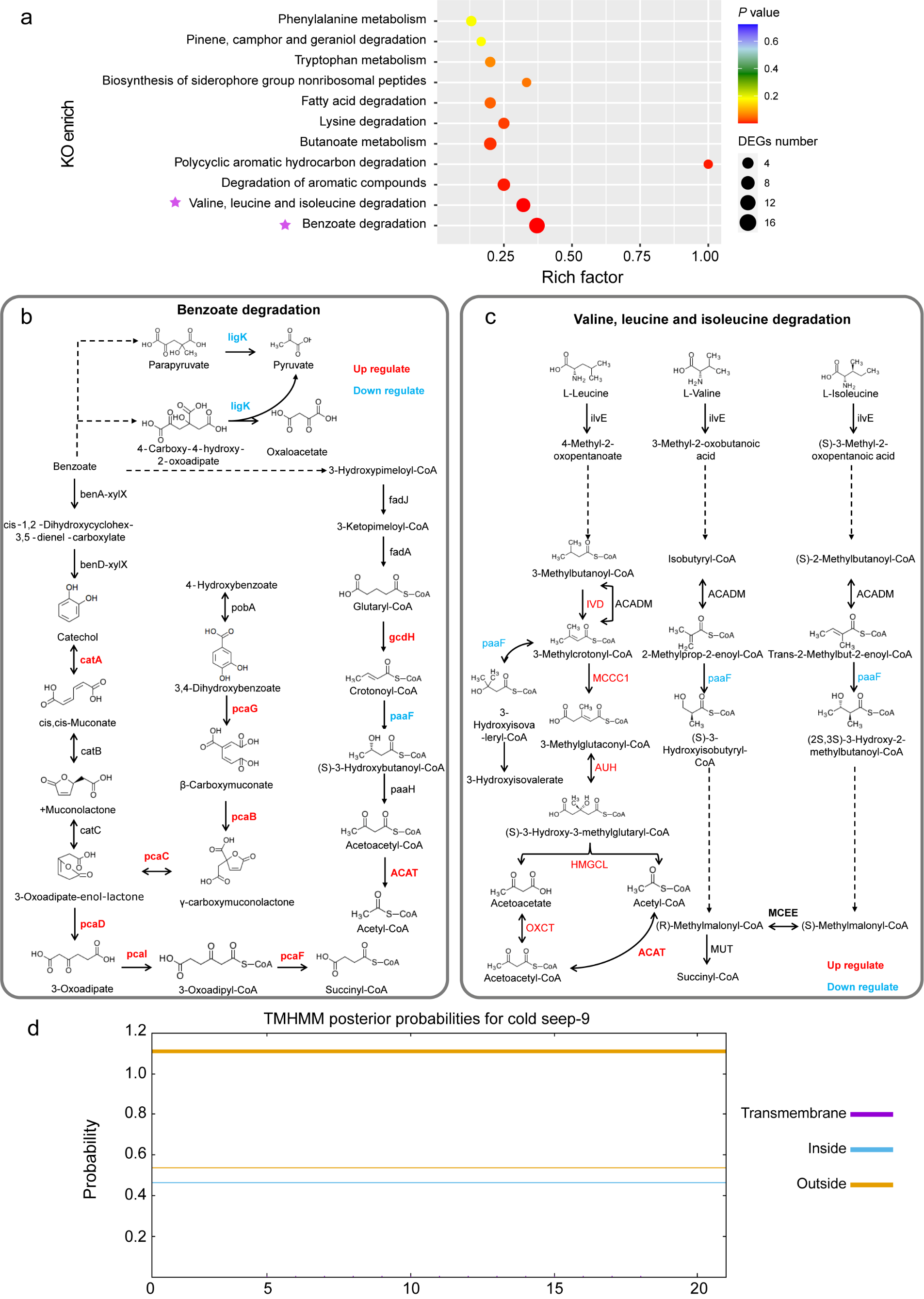

**Supplementary Figure 16. Metabolic pathway enrichment and functional maps of *Acinetobacter pittii* treated with the cold seep-9 antimicrobial peptide. (a)** KEGG pathway enrichment analysis of differentially expressed genes (DEGs), showing enriched pathways (y-axis), rich factor (x-axis), DEG counts (bubble size), and statistical significance (color scale). Pathways marked with stars denote major AMP-affected metabolic routes, including valine/leucine/isoleucine degradation and benzoate degradation. **(b)** Benzoate degradation pathway map, with differentially expressed genes highlighted. Red labels indicate significantly upregulated genes, and blue labels indicate significantly downregulated genes. **(c)** Valine, leucine, and isoleucine degradation pathway, illustrating key enzymatic steps and DEGs. Upregulated and downregulated genes are indicated in red and blue, respectively. **(d)** TMHMM topology prediction for peptide cold seep-9, showing posterior probabilities for transmembrane, intracellular, and extracellular regions across its amino acid sequence.

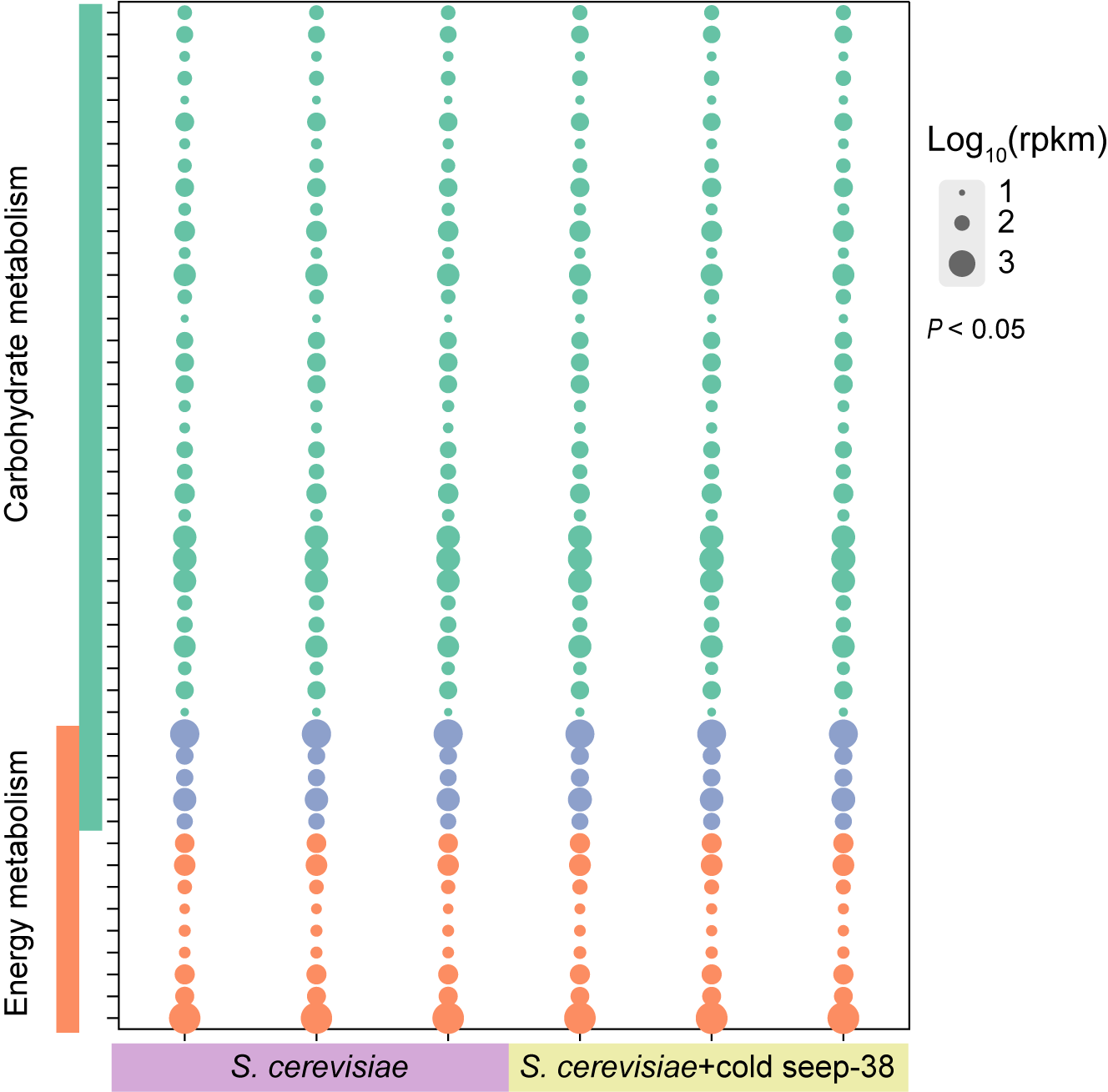

**Supplementary Figure 17. Bubble plot showing transcriptomic responses of *Saccharomyces cerevisiae* treated with the cold seep-38 antimicrobial peptide.** Expression levels of genes associated with carbohydrate metabolism and energy metabolism in *S. cerevisiae* (left) and *S. cerevisiae* treated with cold seep-38 (right). Bubble size represents gene expression level (log_10_ (RPKM)), and only genes with significant differential expression (*P* < 0.05) are shown. The color of each bubble indicates the metabolic pathways in which the corresponding genes are involved: orange represents genes involved in energy metabolism, green represents genes involved in carbohydrate metabolism, and blue represents genes involved in both energy and carbohydrate metabolism.

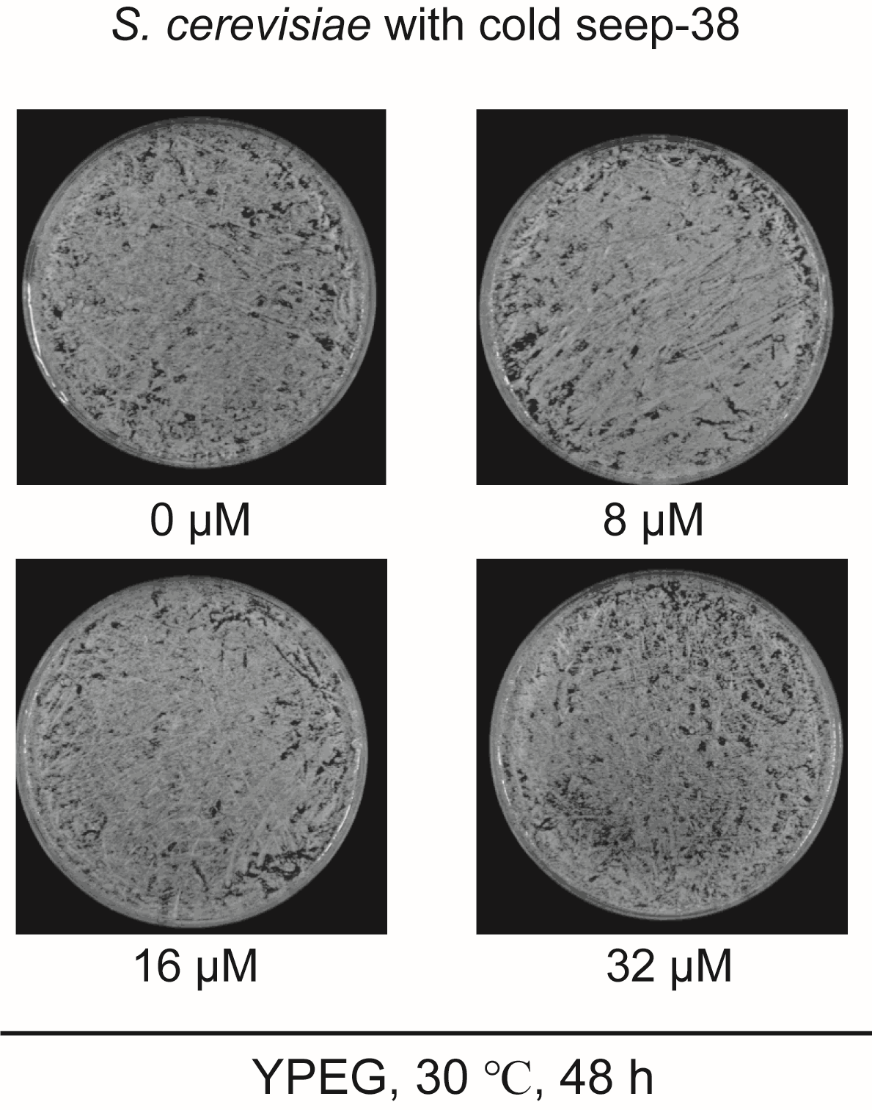

**Supplementary Figure 18. Post-treatment growth of *Saccharomyces cerevisiae* following removal of cold seep-38 after exposure to different concentrations of the antimicrobial peptide.** *S. cerevisiae* cultures were exposed to cold seep-38 at 0, 8, 16, and 32 μM, after which the peptide was removed and cells were plated on YPGE medium, and incubated at 30 °C for 48 h. Representative plates show colony formation under each condition. After removal of cold seep-38, cells recovered and displayed normal growth on YPGE, indicating that the inhibitory effect of the peptide is reversible.

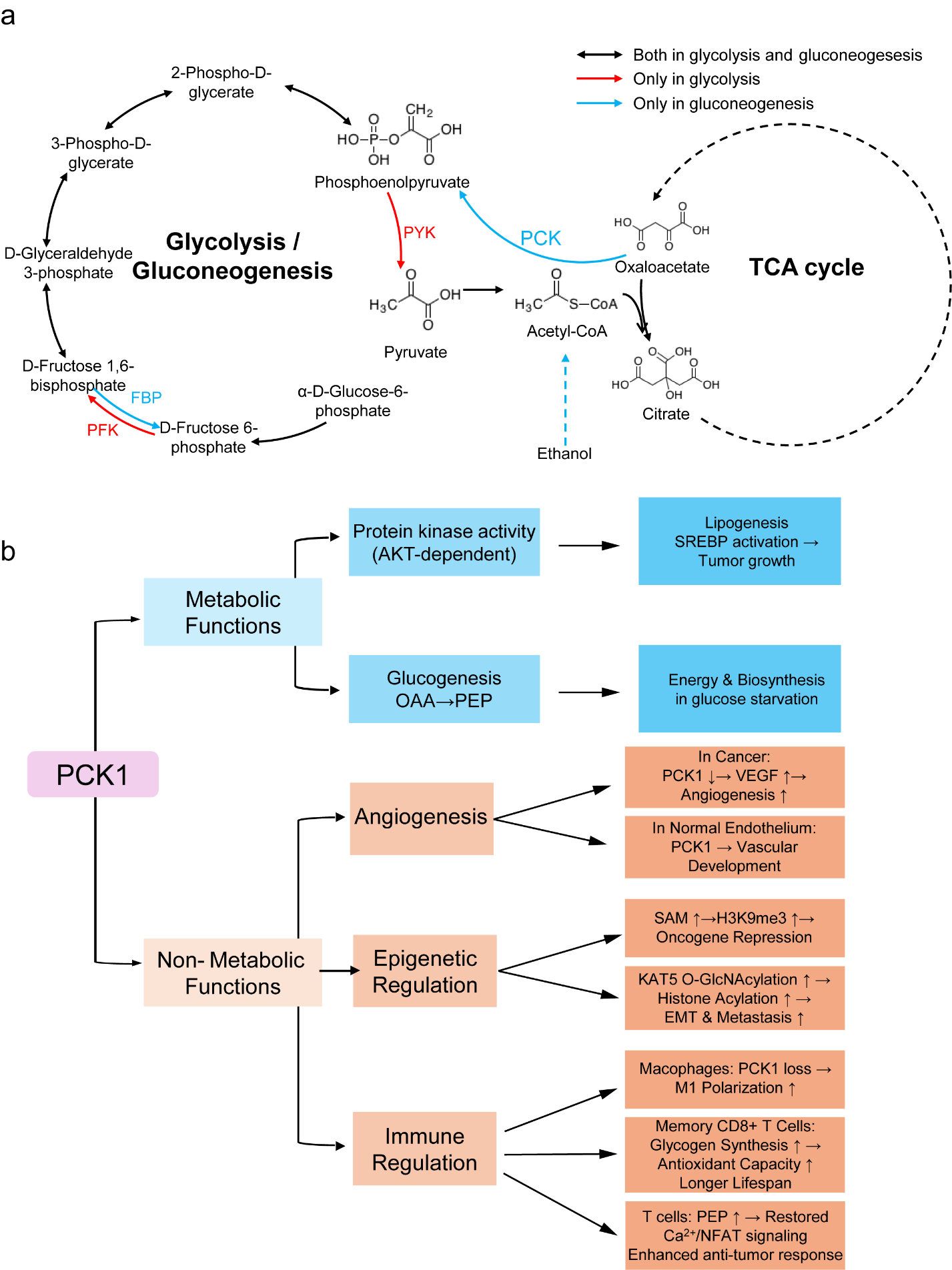

**Supplementary Figure 19. PCK1 as a central hub linking YPGE metabolism to tumor and immune functions. (a)** YPGE pathway overview: PCK1 (blue) catalyzes the rate-limiting gluconeogenic step (oxaloacetate → phosphoenolpyruvate) using TCA cycle intermediates. Red and blue arrows indicate glycolysis- or gluconeogenesis-specific reactions, while black arrows denote shared steps. **(b)** PCK1 functions beyond metabolism: Metabolically, PCK1 promotes lipogenesis and tumor growth via AKT-dependent protein kinase activity and supports energy production under glucose starvation through gluconeogenesis. Non-metabolically, PCK1 regulates tumor angiogenesis (VEGF signaling), remodels epigenetic states (histone modifications), and shapes immune responses, including macrophage polarization and T-cell survival and anti-tumor activity.
